# Glycan Atlassing: Nanoscale analysis of glycocalyx architecture enables functional tracing of cell state

**DOI:** 10.1101/2025.04.29.651200

**Authors:** Dijo Moonnukandathil Joseph, Nazlican Yurekli, Sarah Fritsche, Reem Hashem, Oana-Maria Thoma, Imen Larafa, Tina Boric, Chloé Bielawski, Karim Almahayni, Kristian Franze, Maximilian J. Waldner, Leonhard Möckl

## Abstract

The glycocalyx is a complex layer of glycosylated biomolecules surrounding all cells in the human body. It is involved in the regulation of critical cellular processes such as immune response modulation, cell adhesion, and host-pathogen interactions. Despite these insights, the functional relationship between glycocalyx architecture and cellular state has remained elusive so far, mainly attributable to the structural diversity of glycocalyx constituents and their nanoscale organization. Here, we show that DNA-tagged lectin labeling and metabolic oligosaccharide engineering enables multiplexed super-resolution microscopy of glycocalyx constituents, yielding an atlas of glycocalyx architecture with nanometer resolution. Quantitative analysis of the obtained nanoscale map of glycocalyx constituents facilitates the extraction of characteristic spatial relationships that accurately report on cellular state. We demonstrate the capacity of our approach, which we term Glycan Atlassing, across cell and tissue types, ranging from cultured cell lines to primary immune cells, neurons, and primary patient tissue. Glycan Atlassing establishes a powerful strategy for investigating glycocalyx remodeling in development and disease, potentially enabling the development of new glycocalyx-centered targets in diagnosis and therapy.

## Introduction

The glycocalyx is a dense and dynamic layer of glycosylated molecules that surrounds all mammalian cells, making it the first part of the cell to interact with the exterior (Figure 1A).^1,2^ It has been shown that the glycocalyx is involved in various cellular processes. These processes include, but are not limited to, immune response modulation,^3^ blastocyst adhesion,^4,5^ and host-pathogen interactions.^6^

**Figure 1:**
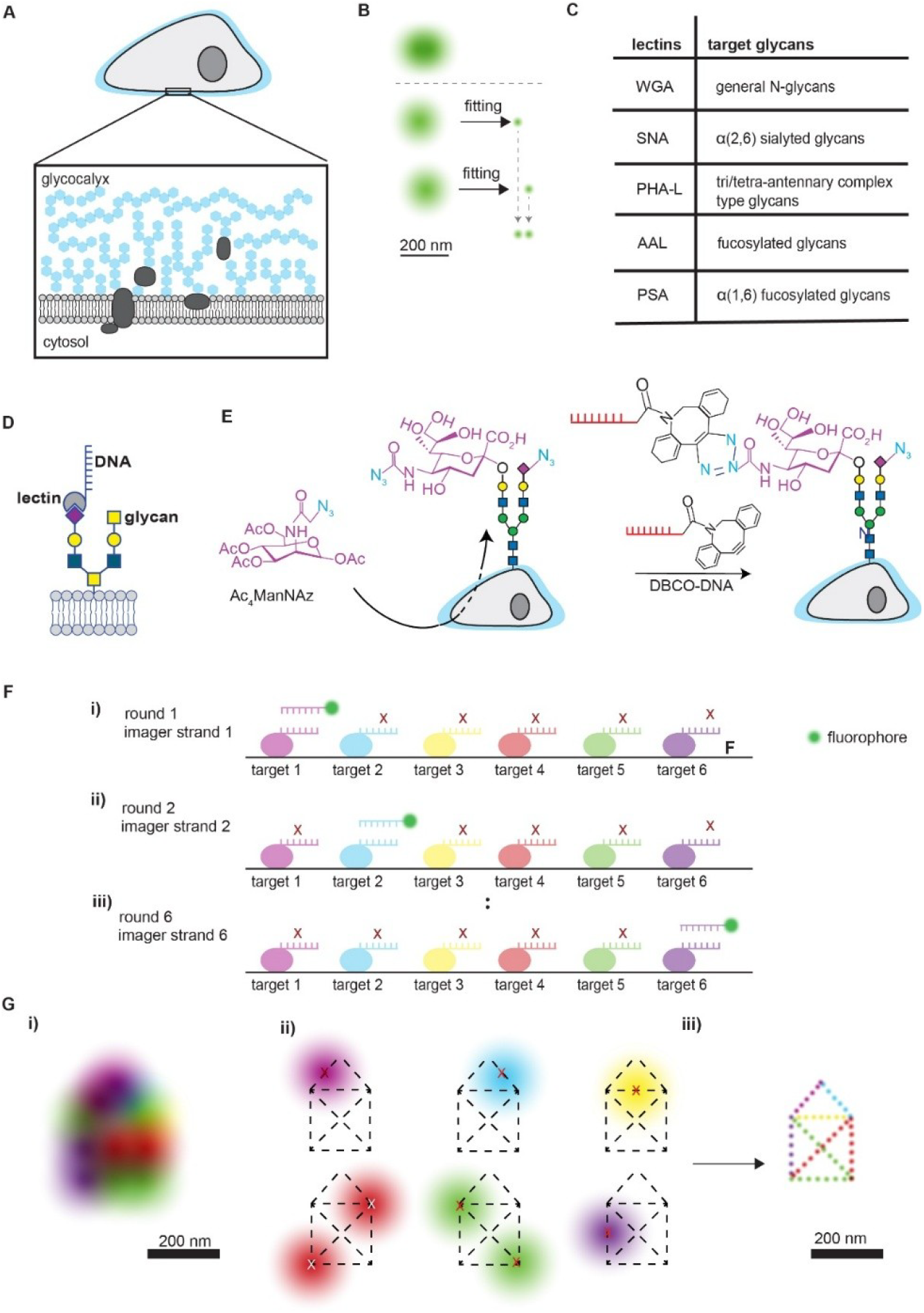
Glycan Atlassing reveals the nanoscale organization of the glycocalyx. **A)** Schematic representation of the glycocalyx. **B)** Working principle of single-molecule localization microscopy. Separation of fluorophores spaced closer than the diffraction limit in time. **C)** Table of lectins employed in this study and their target glycan structure. **D)** Introduction of a DNA barcode via labeling with tagged lectins. **E)** Schematic overview of metabolic labeling with Ac_4_ManNAz, targeting sialic acids. **F)** Six-plex super-resolution imaging enabled by tagging six different imaging targets with six orthogonal DNA strands. (i) to (iii) depict three out of six imaging rounds. In each round, only one target is imaged due to the orthogonality of the DNA strands. **G)** A structure with features below the diffraction limit cannot be resolved with conventional light microscopy (i). Sequential imaging (ii) enables resolution of sub-diffraction features (iii).

In the past decades, studies have revealed numerous aspects of the functional role of the glycocalyx in cell biology. For example, interactions between sialylated ligands and immune cell siglecs were shown to reduce immune cell activity,^7^ an increase of glycocalyx thickness was observed upon oncogenic events,^8^ and the interplay between altered glycosylation and metastasis has been investigated.^9-11^

From a structural point of view, a key component of the glycocalyx are glycans, which consist of monosaccharides such as glucose, galactose, and sialic acids. The structural diversity of glycans found within the glycocalyx is immense due to their various sizes, molecular compositions, and stereochemical properties.^12^

A relationship between the highly complex structural organization and the biological function of the glycocalyx in health and disease seems likely, but is, to date, not comprehensively understood. Analytical methods, such as mass spectrometry, are powerful in identifying glycan structures, but cannot infer details on the native spatial arrangement.^13^ Mass spectrometry imaging partially addresses this drawback by ionizing glycans from the native cellular environment, however, its resolution does not allow for subcellular, not to mention molecular-scale analyses.^14^ Furthermore, electron microscopy has been used to study the glycocalyx.^15^ Although it offers high resolution, concerns persist that its sample preparation procedures might harm the native glycocalyx structure. Moreover, electron microscopy provides only grayscale images of electron density, lacking the species contrast needed to distinguish between molecular components.

Optical microscopy is considered comparably non-invasive and provides species contrast.^16^ However, the nanoscale architecture of the glycocalyx precludes the use of conventional optical microscopy as its resolution is limited by diffraction to approx. 250 nm.^17^ This changed with the advent of super-resolution microscopy methods. In particular, single-molecule super-resolution microscopy techniques rely on the temporal separation of fluorophore emission. If two fluorophores spaced closer than the diffraction limit emit at the same time, their diffraction-limited intensity profiles overlap on the camera, precluding their separation (Figure 1B). A control mechanism (either targeted or stochastic) can be applied to force them to emit at different times. In this case, the individual intensity profiles can be fitted with an appropriate model function (e.g., a 2D Gaussian), which enables the localization of the emitter position with high accuracy, well below the diffraction limit.^18-20^

This approach has facilitated the first analyses of glycocalyx constituents at sub-diffraction length scales.^8,21,22^ Recent work has employed multiplexed dSTORM to analyze the relative abundance of lectin-labeled glycans in different cellular organelles.^23^ Additionally, lectin-based staining of several glycocalyx constituents was employed to analyze diffusion metrics in service of extracting cellular parameters from the motion of cell-surface glycans.^24^ Taken together, previous studies have impressively demonstrated the potential of super-resolution microscopy techniques for glycocalyx analysis.

Despite these advances, comprehensive quantification of glycocalyx architecture at the nanoscale and its connection to functional cell state has not been achieved yet. Here, we present Glycan Atlassing, a rigorous quantitative mapping of cell-surface glycan architecture. We employ lectin labeling in combination with metabolic incorporation of unnatural sugars in order to tag distinct glycocalyx constituents with DNA sequences optimized for high-speed multiplexed super-resolution microscopy.^25^ The obtained localization data is subsequently analyzed via a tailored pipeline that extracts characteristic spatial relationships within glycan moieties, both at the level of individual and grouped localizations (software package GlycanConstructor^26^ or “GlyCo”). With this approach, we, for the first time, establish a super-resolved atlas of the glycocalyx, connecting nanoscale glycocalyx organization and cellular state across a range of sample types – from cultured cell lines over primary immune cells and neurons to patient tissue.

## Results and Discussion

### Strategy for multiplexed DNA-PAINT microscopy of glycocalyx constituents

Glycan Atlassing centrally employs DNA points accumulation for imaging in nanoscale topography (DNA-PAINT)^27^ with labeling via DNA-barcoded lectins and metabolic oligosaccharide engineering.^28^ With this integrated approach, Glycan Atlassing facilitates super-resolution imaging of glycocalyx constituents (Figure 1C/D). The selection of suitable lectins was based on previous studies investigating lectin performance metrics.^29^ Thus, we obtained a comprehensive overview of commercially available lectins and their binding specificities as well as their affinities and mutual compatibility. After initial screening of the database, a refinement was performed using literature reporting on further experimental studies.^30-32^

Ultimately, we identified the lectins shown in Figure 1C as suitable candidates for subsequent analysis. Wheat Germ Agglutinin (WGA) binds to sialic acid and N-acetylglucosamine, which are upregulated in cancer cells to promote immune evasion and metastasis.^10^ Sambucus Nigra Agglutinin (SNA) identifies α-2,6-linked sialic acids, a modification that correlates with tumor aggressiveness and immune regulation.^33,34^ Phaseolus Vulgaris Leucoagglutinin (PHAL) detects β1-6 branched N-glycans, a hallmark indicative of malignant transformation that enhances tumor invasiveness.^35^ Aleuria Aurantia Lectin (AAL) binds to fucose linked (α-1,3, α-1,2, α-,4, and α-1,6) to N-acetyllactosamine, which play roles in immune recognition and inflammatory processes.^36^ Pisum Sativum Agglutinin (PSA) targets α-fucose residues, involved in pathogen recognition and immune signaling.^37^

Simultaneously, metabolic oligosaccharide engineering was applied by incorporating N-azidoacetylmannosamine (Ac_4_ManNAz) into cell-surface sialic acid residues (Figure 1E). This process introduces azido-modified sialic acids into the glycocalyx through the cell’s native metabolism, allowing for subsequent bioorthogonal reactions.^28^ Following the incorporation of the azido-modified sugar analogue, the metabolically labeled sialic acids was conjugated with DNA-barcoded dibenzocyclooctyne (DBCO) via copper-free click chemistry. The strain-promoted azide-alkyne cycloaddition reaction proceeds rapidly under physiological conditions, forming stable triazole linkages between the azido-modified glycans and DBCO without the need for potentially cytotoxic copper catalysis.^38^

Multiplexed DNA-PAINT microscopy was performed via sequential introduction of six distinct imager strand, each designed to hybridize with a single complementary docking strand without cross-affinity.^25^ Thus, each imager strand exclusively hybridizes with its complementary docking strand. All other targets labeled with different docking strands are not recognized (Figure 1F). After each imaging round, through washing ensured the absence of cross-talk between sequential imaging rounds, and the next imager strand was added. This cycle of imager strand addition, data acquisition, and washing was repeated until all targets were addressed. This process is schematically depicted in Figure 1G. The image achievable in conventional fluorescence microscopy is shown in (i), where details below the diffraction limit are blurred. Sequential acquisition of single-molecule signals for each imaging target and their precise localization (ii) enables the super-resolved reconstruction of structural features (iii).

### Glycan Atlassing enables multiplexed nanoscale analysis of glycocalyx organization across cells and tissues

Glycan Atlassing was established on a range of relevant sample types with increasing complexity, ensuring that benchmarking of the approach was performed in a controlled environment before moving to more demanding samples. The different systems studied are schematically represented in Figure 2A. We began with standard cultured cell lines (Figure 2B), followed by primary rat neurons (Figure 2C), human primary immune cells (Figure 2D), and finally primary human tissue (Figure 2E). Each sample was labeled and analyzed using the same protocol, which was only slightly adapted to satisfy specific requirements of the systems studied (see Methods). Additional examples are shown in Figure S1.

**Figure 2:**
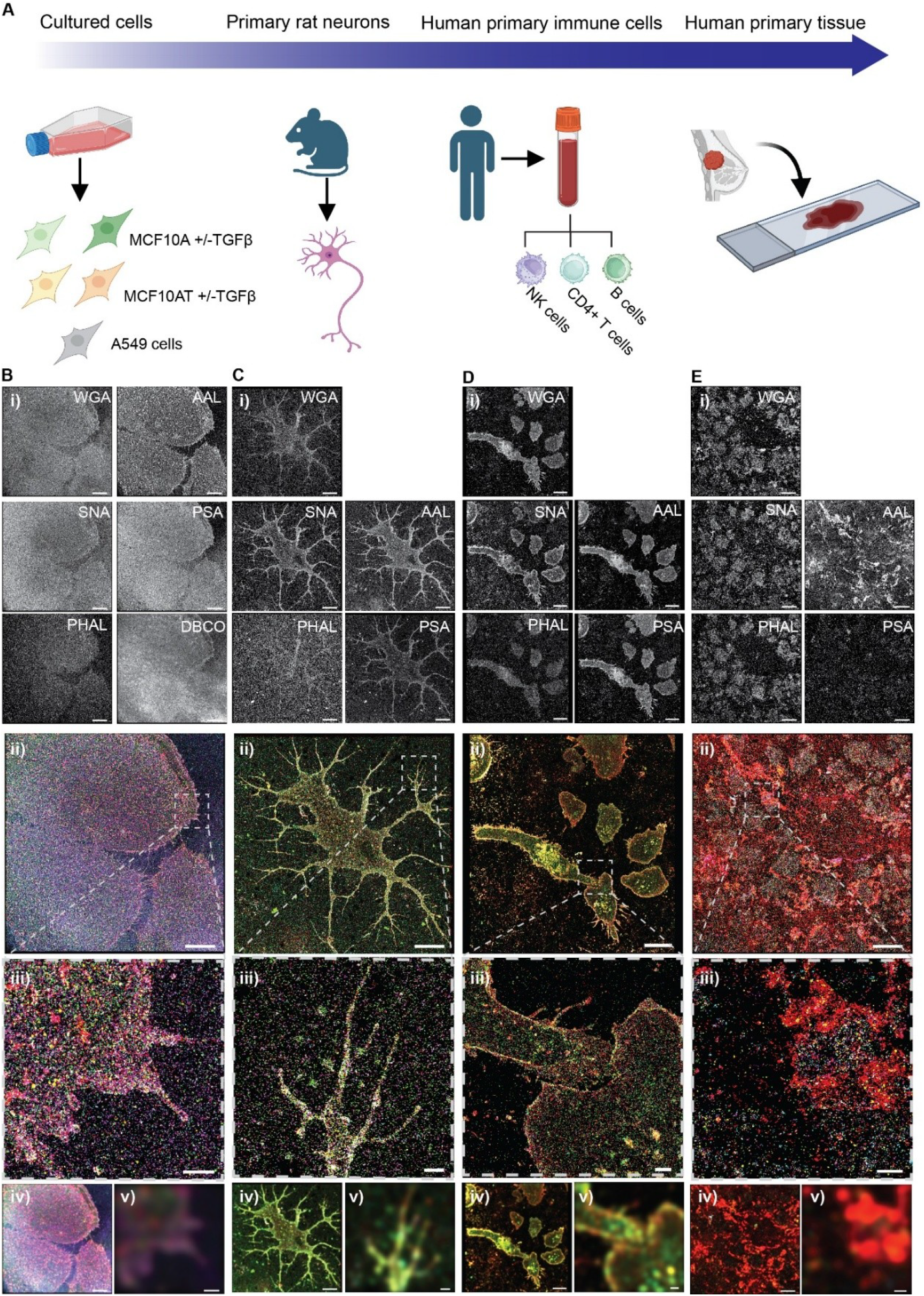
Overview of investigated systems. **A)** Sample library investigated in this study. **B-E)** Individual channels depicted in grayscale (i), merged channels (ii) using the following color code: WGA – magenta, SNA – cyan, PHA-L – yellow, AAL – red, PSA – green, DBCO – purple. iii) Zoom-in showing intricate details resolved. iv) Diffraction-limited representation of the whole field of view. v) Diffraction-limited zoom-in corresponding to (iii). Scale bars: 10 µm for full field of views; 1 µm for zoom-ins.

In brief, the samples were incubated with DNA-barcoded lectins. For MCF10A/AT cells, they were also incubated with Ac4ManNAz for metabolic labeling of sialic acids, followed by copper-free click chemistry to attach the docking strand. The samples were then subjected to multiplexed DNA-PAINT microscopy, data analysis, and quantitative spatial mapping of the obtained high-dimensional nanoscale imaging data.

With this approach, we gradually moved from *in vitro* to *ex vivo*, achieving a robust demonstration of Glycan Atlassing across cells and tissues. For each sample type shown in Figure 2B-E, reconstructions from individual imaging rounds targeting different glycan species are presented in grayscale alongside the respective labeling agent (i). Merged reconstructions of the full field of view (ii) and magnifications (iii) are shown alongside the diffraction-limited equivalents (iv, v), highlighting the substantial gain in resolution and the successful visualization of nanoscale glycocalyx structure. Notably, differences between lectins are evident at the qualitative level depicted here, as well as differences between identical lectins on different sample types. This indicates that our approach has target- and sample-specific readout potential, even at the qualitative level, which we extend to the quantitative level as described below.

### Quantitative analysis pipeline for extraction of characteristic glycan patterns

Due to the structural intricacy of the glycocalyx, the data obtained by super-resolution investigation of its architecture is inherently complex. To decipher this complexity, it is not sufficient to merely visually inspect the reconstructions. Thus, Glycan Atlassing employs quantitative analysis of spatial signatures, obtained by the investigation of nanoscale relationships between glycan species on the cell surface.

Figure 3A shows the key steps in Glycan Atlassing. Raw localization data is captured sequentially for each imaging target. This localization data is drift corrected and localizations for each imaging target are aligned (both using gold nanoparticles as fiducials). This is followed by segmentation of the region of interest. Then, clustering is performed to group multiple localizations arising from the repeated interaction between imager strands and a single docking strand. The cluster centers are taken as the target location, which we will call “lectin binding sites” in the following discussions. Subsequently, quantitative analysis using nearest neighbor peak distance matrices and grouped target locations using GlycCo is performed, followed by dimensionality reduction via principal component analysis (PCA).

**Figure 3:**
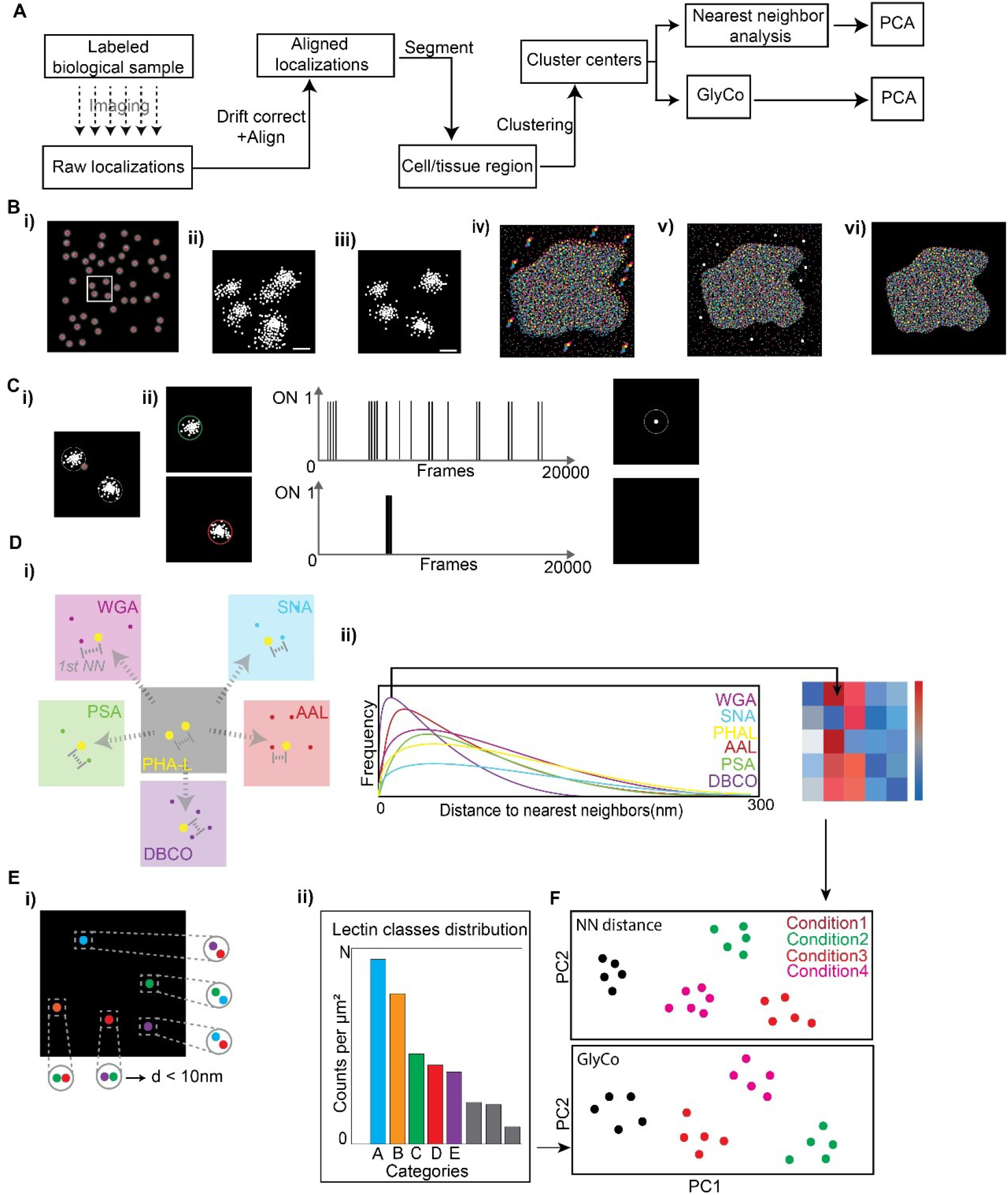
Glycan Atlassing data acquisition and analysis pipeline. **A)** Experimental flow of Glycan Atlassing. **B)** Central steps of data processing. i) Single-molecule signals are localized. ii) Multiple samplings of a single docking strand results in localization “clouds” (here, four docking strands are depicted corresponding to the white box in (i)). iii) Drift correction. iv) All channels overlaid before alignment. v) Alignment via gold particles as spatial fiducial markers. vi) Segmentation of the target cell. **C)** Separation of true imager-docking strand interactions and unspecific sticking events. i) Two exemplary localization clouds. ii) Characterization of the time traces of the interactions. Top: True imager-docking strand interactions are short-lived and repeat. Bottom: Unspecific sticking events do not repeat and are long-lived. Sticking events are filtered out. **D)** Principle of nearest neighbor (NN) analysis i) The distance between every target location and its nearest neighbor in the same and in all other channels are calculated. ii) Nearest neighbor histograms of one representative channel. ii) Color-coded matrix reporting on all inter- and intra-channel peaks of NN distance histograms. **E)** Principle of GlyCo. Two or more target locations from different channels are grouped if their distance is smaller than a threshold. i) Five example classes, each containing two target locations, are shown. ii) The ten most frequent classes found in a sample are collected. **F)** Dimensionality reduction. Principal component analysis (PCA) is conducted on the NN distance matrices and the histograms from GlyCo.

In detail, single-molecule emissions are captured as an image stack of 20,000 frames within each imaging round (Figure 3B). The precise localization of each individual emitter position is determined with nanoscale accuracy via fitting with a two-dimensional Gaussian (i). Figure S2 shows all relevant quality metrics for the fitting procedure across dataset in this study. In DNA-PAINT, each docking strand is addressed by complementary imager strands multiple times. Each interaction yields a separate localization. These localizations are spread to a “cloud” of localizations around the true position of the docking strand due to experimental factors such as slightly different camera noise or background fluorescence (ii). Furthermore, thermal instabilities cause the sample to drift over time, resulting in the localizations to be seen as a smeared cloud. This drift is accounted for by tracking the apparent movement of immobile gold fiducial beads added to the sample before imaging. By extracting the apparent movement, the drift can be removed (iii). Localizations from all the imaging targets create separate nanoscale spatial distributions of each target glycan (corresponding to one channel). These channels are merged (iv) and aligned with sub-nanometer precision by employing gold fiducial beads (v). For further processing, manual segmentation of the region of interest is performed.

In DNA-PAINT, a potential source of false positive localizations arises from unspecific sticking events, i.e., an imager strand binding to some cellular structure, causing a localization that is not arising from an interaction with a docking strand. These sticking events are rare; however, they need to be corrected for in order to minimize perturbation of subsequent analyses (Figure 3C). Sticking events can be detected by abnormal interaction time signatures of imager strands. Compared to short-repetitive docking-imager interactions (i), sticking events are usually longer-lived and non-repetitive (ii). Thus, by characterizing the time signatures of imager strand interactions, sticking events are efficiently removed.

Ultimately, multiplexed spatial distributions of glycocalyx constituents is obtained with nanometer resolution. Further analysis extracts the relationships between the imaging targets. We implemented two fundamental types of analysis. First, we determine the NN distance distributions within one channel as well as between different channels (Figure 3D). Nearest neighbor distances are plotted (i). The most frequent NN distances from all comparisons are collected (ii). Second, we identified glycan species who are frequently closely related, classifying them according to their simultaneous occurrence in space. For this, we developed Glycan Constructor (GlyCo) (Figure 3E), which groups localizations across channels within a radius of 5 nm (i) and plots the distribution of observed classes per unit area (ii) along with a location map of the different classes. We note that the threshold of 5 nm can be set differently, however, considering that the typical sizes of individual glycans usually range between 5 to 10 nm, this setting is reasonable.^39^ These two results – NN peak distance matrices and classes of frequently related target locations – are then subjected to dimensionality reduction using a standard PCA, projecting the high-dimensional input data to three dimensions. (Figure 3F).

### Glycan Atlassing enables tracing of cancer progression

As a first imaging target, we turned to immortalized mammary epithelial cells (MCF10A cells). In addition, a transformed variant (MCF10AT), constitutively expressing the oncogene HRAS was included. Both cell lines were either treated with the tumor growth factor β (TGFβ) or left untreated. Thus, we investigated the panel of MCF10A, MCF10A+TGFβ, MCF10AT, MCF10AT+TGFβ as a model system for epithelial-to-mesenchymal transition (EMT), a key step in cancer progression.^40,41^ In combination with lectin labeling, we performed metabolic labeling of terminal sialic acids on these cells.

Figure 4A shows localization data from a representative MCF10A cell. Raw localizations from individual rounds of imaging are shown in grayscale (i) followed by the merged localizations (ii) and the brightfield image (iii). An arrowhead points towards the cell whose analysis is shown in the Figure. Figure 4B shows an example region where clustering of raw localizations yields the lectin binding sites. NN distributions for a representative channel, here PHA-L, are shown in Figure 4C, which report on the distance distribution of neighbors for complex N-glycans (the labeling target of PHA-L). The peak location of the NN distributions for glycan shown in the matrix in Figure 4D. The distribution of lectin classes from classifying spatially related target locations using GlyCo are shown in Figure 4E alongside their arrangement on the cell surface in Figure 4F. Additional data is shown in Figure S3.

**Figure 4:**
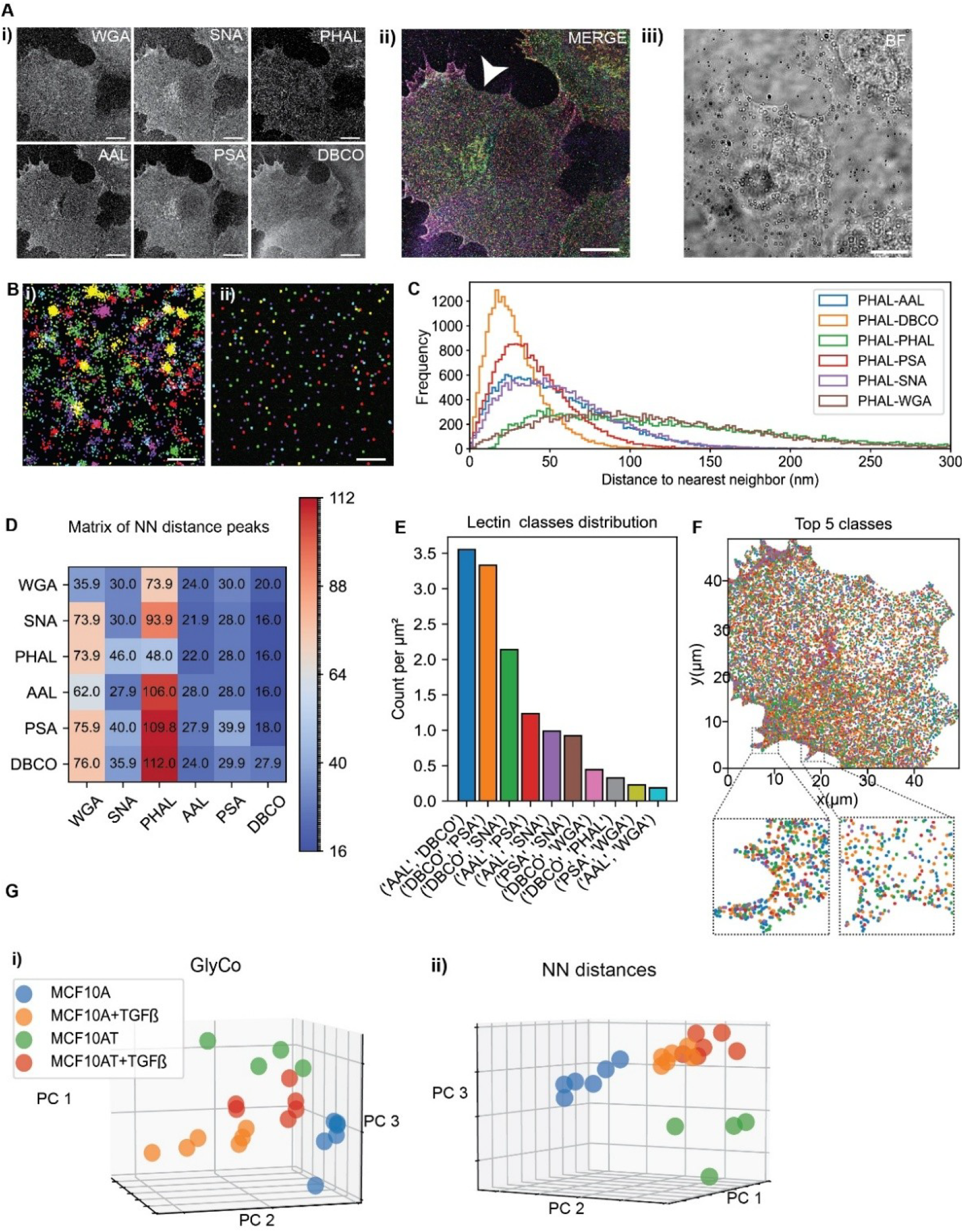
Glycan Atlassing traces early oncogenic events. **A)** i) Individual channels in grayscale ii) Merge of individual channels (the white arrowhead indicates the cell analyzed here; see Figure 2 for color map). iii) FOV in bright field. **B)** Representative example of clustering localization clouds to single target locations i) Raw localizations ii) Clustered localizations. **C)** NN histograms of one representative channel (here: relationship of PHA-L to all channels) **D)** Maxima from NN histograms across all comparisons. **E)** Lectin Class distribution from GlyCo. Ten most frequent classes and their spatial density are shown. **F)** Spatial arrangement of the classes shown in E). **G)** PCA plots for i) GlyCo and ii) NN peak distances for all samples investigated, comparing the four stages of EMT modeled. Scale bars: 10 µm in A), 100 nm in B).

Several intriguing observations can be made from the data. First, qualitative differences in local densities within one imaging target as well as global differences between imaging targets are evident. Furthermore, spatial distributions of different sialic acid and fucose linkages can be observed at nanometer resolution. Finally, and most strikingly, PCA (Figure 4G) of the multidimensional datasets from GlyCo (i) and NN peak distribution (ii) is able to faithfully separate the different stages of cancer progression within the MCF10A-based cellular model system. It is worth pointing out that the clusters of TGFβ-stimulated cells are located closer to each other than the unstimulated conditions. This indicates that TGFβ stimulation drives both MCF10A and MCF10AT cells to a closely related hyperproliferative phenotype, which is mirrored in the glycocalyx state. Taken together, the characteristic nanoscale glycocalyx signatures accurately report on the internal cellular state, and Glycan Atlassing is able to retrieve this information.

### Glycan Atlassing enables analysis of subcellular neuronal glycosylation

Next, we proceeded to employ Glycan Atlassing on embryonic rat hippocampal neurons. Primary neurons were extracted from embryonic rat hippocampal tissue (see Methods for details). The cells were cultured on glass-bottomed petri dishes coated with PDL and laminin for 5 to 6 days, and subsequently fixed in 4% PFA.

Figure 5 shows the results of Glycan Atlassing on primary rat hippocampal neurons, following the structure of Figure 4. Expression of individual glycan species within the neuronal glycocalyx are shown in Figure 5A. Figure 5B depicts raw localizations and clustering to obtain individual lectin binding sites. NN distance distributions as well as the full NN distance peaks are shown in Figure 5C/D. Classification of spatially related target glycans using GlyCo is shown in Figure 5E/F. In this case, we investigated differences in the nanoscale glycosylation patterns between the cell body and dendrons of neurons. Finally, the analysis of all investigated samples using PCA-based dimensionality reduction is reported in Figure 5G. Additional data is shown in Figure S4.

**Figure 5:**
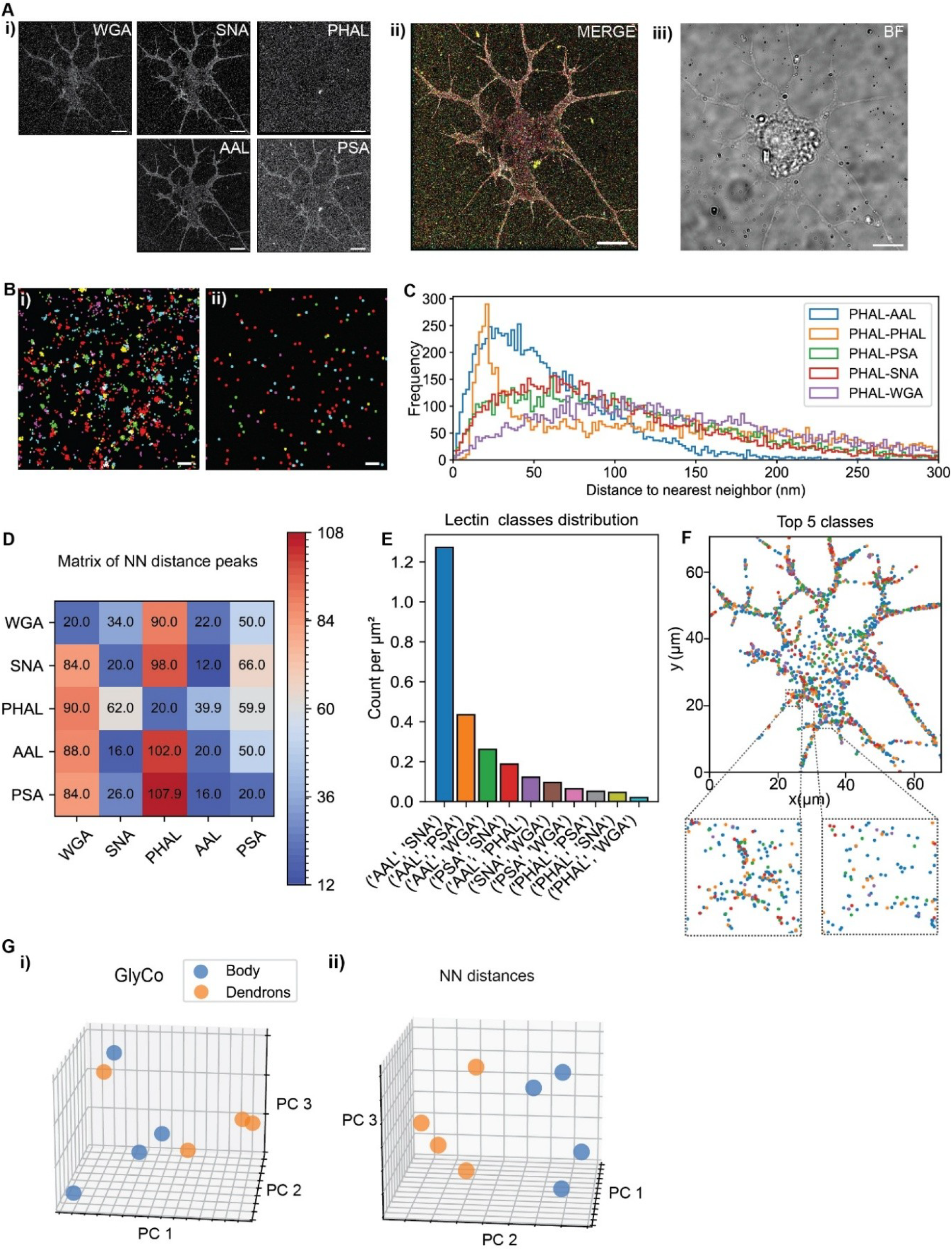
Glycan Atlassing reveals characteristic neuronal glycosylation patterns. **A)** i) Individual channels in grayscale ii) Merge of individual channels (see Figure 2 for color map). iii) FOV in bright field. **B)** Representative example of clustering localization clouds to single target locations i) Raw localizations ii) Clustered localizations. **C)** NN histograms of one representative channel (here: relationship of PHA-L to all channels) **D)** Maxima from NN histograms across all comparisons. **E)** Lectin Class distribution from GlyCo. Ten most frequent classes and their spatial density are shown. **F)** Spatial arrangement of the classes shown in E). **G)** PCA plots for i) GlyCo and ii) NN peak distances for all samples investigated, comparing characteristic glycosylation patterns between the neuron cell body and dendrons. Scale bars: 10 µm in A), 100 nm in B).

Interestingly, PHA-L signal is low in the neurons investigated, consistent with the reported timing of complex N-glycosylation (the labeling target of PHA-L) in primary neuronal culture. This observation thus supports our analysis, as elaborate N-glycosylation is expected to become prominent in matured primary neurons, whereas the cells investigated here are considered in an early stage of development.^42^

As before, Glycan Atlassing is able to distinguish between the analyzed conditions based on characteristic glycan patterns. The PCA yields comparably less clustered data than for the analysis of the EMT panel (Figure 4), however, this is expected, given that we compare different parts of identical cells, which are expected to be similar.

Overall, Glycan Atlassing reveals differential glycosylation signatures within individual neurons, which may indicate an unidentified regulatory axis. For example, glycosylated proteins may engage with galectins, modulating membrane receptor residence times and internalization kinetics as previously reported.^43^

### Glycan Atlassing identifies immune cell state

To investigate the functional glycocalyx architecture of immune cells in different cellular states, we next performed Glycan Atlassing on a range of different primary immune cells. In particular, we investigated primary NK cells, primary CD4+ T cells, and primary neutrophils. These cell types were selected due to their central roles in the immune system and their distinct contributions to immune responses.

NK cells were isolated from whole blood via bead-based separation and cultivated for maximally three passages. For activation, NK cells were co-cultured with human lung cancer-derived A549 cells. Secreted proteins and vesicles from A549 rapidly stimulated NK cells, as evident by their directed movement towards A549 cells. Non-activated NK cells were cultured in isolation. CD4+ T-cells were isolated from whole blood of healthy donors using commercially available kits. After isolation, CD4+ T-cells were in vitro activated using α-CD3/α-CD28 in combination with IL-2 for three days. As control, freshly isolated CD4+ T-cells from same donor were used. Similarly, neutrophils were isolated from whole blood of healthy donors using commercially available kits, according to manufacturer’s instructions. After isolation, the neutrophils were cultured in vitro using human recombinant TNF-α for two hours. Non-stimulated neutrophils were also plated as control (see methods for details)

Figure 6 shows the results of Glycan Atlassing the panel of primary immune cells, following the structure of Figure 4. Expression of individual glycan species within the neuronal glycocalyx are shown in Figure 6A. Figure 6B depicts raw localizations and clustering to obtain individual lectin binding sites. NN distance distributions as well as the full NN distance peaks are shown in Figure 6C/D. Classification of spatially related target glycans using GlyCo is shown in Figure 6E/F. Finally, the analysis of all investigated samples using PCA-based dimensionality reduction is reported in Figure 6G-I, comparing activated and non-activated immune cells. Additional data is shown in Figure S5.

**Figure 6:**
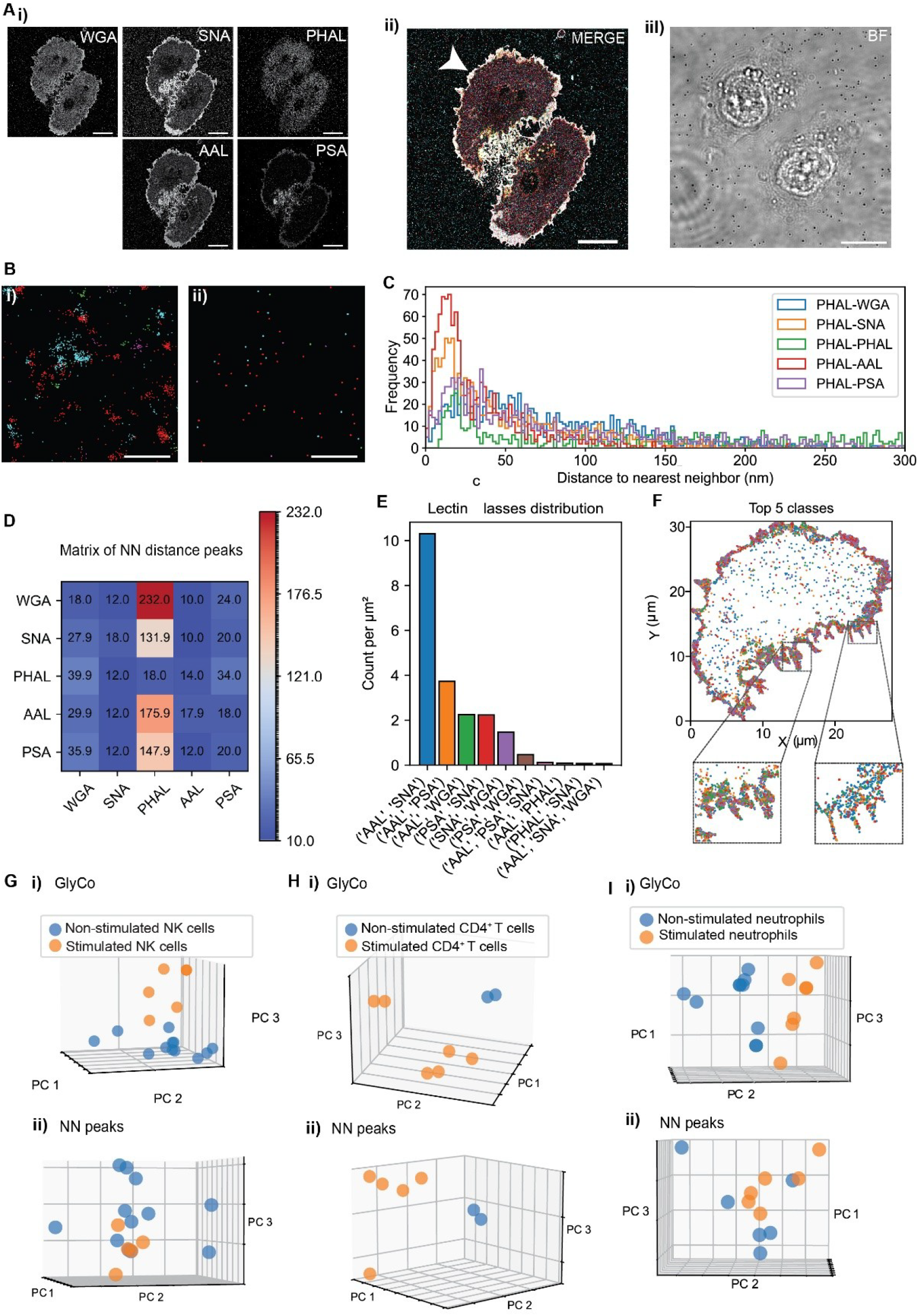
Glycan Atlassing traces immune cell activation. **A)** i) Individual channels in grayscale ii) Merge of individual channels (the white arrowhead indicates the cell analyzed here, see Figure 2 for color map). iii) FOV in bright field. **B)** Representative example of clustering localization clouds to single target locations i) Raw localizations ii) Clustered localizations. **C)** NN histograms of one representative channel (here: relationship of PHA-L to all channels) **D)** Maxima from NN histograms across all comparisons. **E)** Lectin Class distribution from GlyCo. Ten most frequent classes and their spatial density are shown. **F)** Spatial arrangement of the classes shown in E). **G)-I)** PCA plots for i) GlyCo and ii) NN peak distances for all samples investigated, comparing characteristic glycosylation patterns between activated and non-activated NK, CD4+ T-cells, and neutrophiles, respectively. Scale bars: 10 µm in A), 100 nm in B).

Glycan Atlassing enables the precise identification of the activation state of NK cells, CD4+ T-cells, and neutrophils. This achievement is particularly noteworthy given the short timeframe of immune cell activation (e.g., five minutes in case of NK cells) employed in this study. Earlier studies have suggested that the full turnover of the cell-surface glycome occurs within a relatively longer period of 24-48 hours.^44^ However, these studies focused on global turnover of glycocalyx bulk, not on more subtle structural changes.

Our analysis suggests that immune cells are capable of rapid adaptations in their cell-surface glycosylation patterns in response to external stimuli. This indicates that the glycocalyx is remarkably dynamic at the time scale of minutes and highly responsive to the cellular environment, displaying distinct and clearly separatable glycan patterns to the outside. The ability to rapidly modulate cell-surface glycosylation patterns might be a critical mechanism by which immune cells coordinate their responses to pathogens and other external stimuli.

### Glycan Atlassing differentiates healthy and tumor tissues based on characteristic glycan signatures

Finally, we employed Glycan Atlassing on primary patient tissue slices. We investigated primary breast adenocarcinoma tissue, with tumor regions and surrounding non-tumor regions present in the same slice, allowing for internal referencing. The categorization into tumor and non-tumor regions was performed by histopathological grading. Samples were obtained from the central biobank of the University Clinic Erlangen.

Figure 7 shows the results of Glycan Atlassing the panel of primary patient tissue slices, following the structure of Figure 4. Expression of individual glycan species within the neuronal glycocalyx are shown in Figure 7A. Figure 7B depicts raw localizations and clustering to obtain individual lectin binding sites. NN distance distributions as well as the full NN distance peaks are shown in Figure 7C/D. Classification of spatially related target glycans using GlyCo is shown in Figure 7E/F. Finally, the analysis of all investigated samples using PCA-based dimensionality reduction is reported in Figure 7G, comparing tumor and non-tumor regions. Additional data is shown in Figure S6.

**Figure 7:**
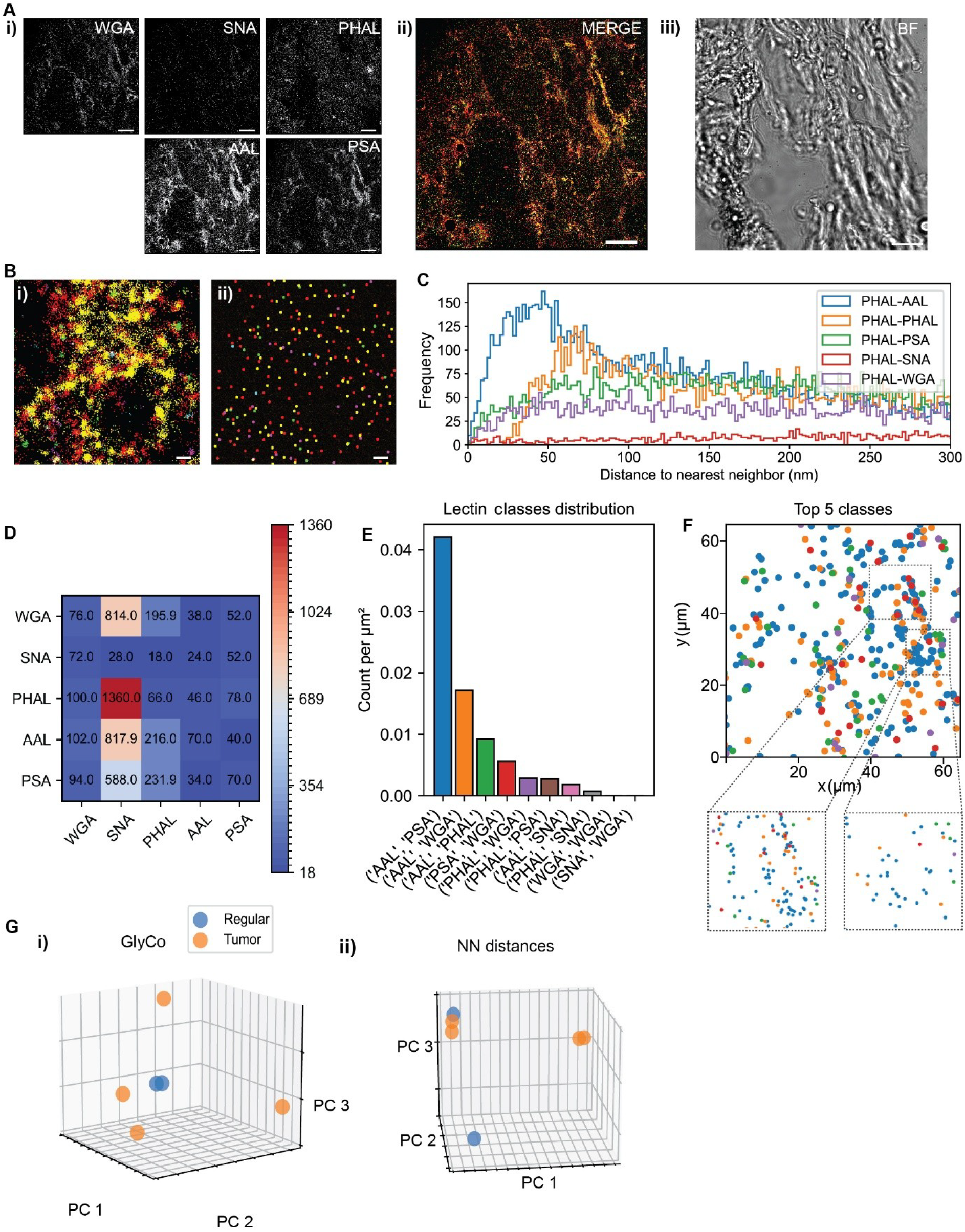
Glycan Atlassing separates healthy and tumor tissue. **A)** i) Individual channels in grayscale ii) Merge of individual channels (the white arrowhead indicates the cell analyzed here, see Figure 2 for color map). iii) FOV in bright field. **B)** Representative example of clustering localization clouds to single target locations i) Raw localizations ii) Clustered localizations. **C)** NN histograms of one representative channel (here: relationship of PHA-L to all channels) **D)** Maxima from NN histograms across all comparisons. **E)** Lectin Class distribution from GlyCo. Ten most frequent classes and their spatial density are shown. **F)** Spatial arrangement of the classes shown in E). **G)** PCA plots for i) GlyCo and ii) NN peak distances for all samples investigated, comparing characteristic glycosylation patterns between tumor and non-tumor areas. Scale bars: 10 µm in A), 100 nm in B).

Indeed, the PCA-based investigation enabled the identification of tumor and non-tumor regions by analysis of their characteristic nanoscale glycan signatures. Interestingly, non-tumor regions form a tight cluster in the PCA, whereas tumor regions are more spread out, suggesting a more heterogenous glycocalyx landscape, consistent with failed regulation of biosynthetic pathways in cancer cells.

Furthermore, we detected elevated signal levels of the lectins AAL and PSA on tumor slices, indicating increased fucosylation. This observation aligns well with previous studies reporting aberrant fucosylation and fucosyltransferases in cancer cells.^45,46^ Going beyond such previous studies that identified bulk changes in tumor cell glycosylation, our approach allows for the direct quantification of glycosylation at the nanoscale level down to distinct glycan linkages. Understanding the precise organization of these glycocalyx structures will likely contribute to a better characterization of cancer cell state and, in a more clinical setting, may help in the development of targeted diagnostic and therapeutic strategies.

## Conclusion and Outlook

Here, we report on Glycan Atlassing, comprehensively enabling the characterization of cellular state by tracing nanoscale glycocalyx architecture. Glycan Atlassing integrates DNA-PAINT, bioorthogonal chemistry and lectin-based glycan targeting as well as quantitative analysis. Beyond establishing the methodology and demonstrating it on a range of relevant biological systems – from cultured cell lines over primary cells to patient tissue –, we identify several so far unknown processes that functionally link glycocalyx state to cell state.

In particular, our results indicate that the process of EMT strongly modulates glycocalyx structure. In the light of earlier studies by our group and others suggesting a functional link between the cancer glycocalyx and cancer progression, Glycan Atlassing will likely be a highly valuable approach to characterize glycocalyx state in fundamental cancer biology.^8,47^ Our results further suggest differential glycosylation states at the subcellular level in primary neurons, raising the question how neuronal activity in health and disease is modulated by the nanoscale architecture of the neuronal glycocalyx. Strikingly, we observed rapid reorganization of specific glycosylation patterns in response to immune cell activation on NK cells, CD4+ T cells, and neutrophils. This result points towards a, to date, uncharacterized axis of dynamic glycocalyx adaptation in immune regulation, with profound implications for glyco-immunology and cell-based therapies. Finally, we were able to take steps towards identifying tumor tissue based on characteristic glycan patterns. These results bear significant impact for clinical use of Glycan Atlassing, e.g., by classifying tumors according to their functional glycocalyx fingerprint. This is especially relevant considering the emerging field of immune cell regulation mediated by the Siglec-sialome axis.^48,49^

Despite these advances, Glycan Atlassing as presented here has limitations. First, a critical barrier that needs to be broken is the limited availability of specific glycan-labeling agents. When compared to the labeling tools available for proteins or genomic sequences, glycan-targeted labeling is currently less developed. Progress in this area is needed and will immediately expand the scope of Glycan Atlassing. Second, our study is currently limited to six imaging targets. Recent reports have demonstrated a larger number of accessible labeling targets,^50^ and the implementation of such strategies will be certainly valuable for future development of Glycan Atlassing. We note, however, that the experimental complexity and the time demand scales with the number of imaged species. Thus, any implementation should balance feasibility with scope. Based on the results presented here, we are confident that the current number of imaging targets enables the extraction of meaningful nanoscale glycocalyx signatures. Third, while Glycan Atlassing is fully compatible with clinical samples, its current implementation requires a high level of expertise and hands-on time from the experimenter, which likely precludes broad clinical adaptation. Here, innovation in data acquisition and analysis automation will be of profound importance. Finally, analysis of NN distances currently only considers peak positions of the extracted distributions, essentially projecting the histogram to a single value that is used as input for further analysis. More advanced approaches could likely extract more refined information from the datasets. However, it is worth to highlight that despite this reduction in dataset complexity, precise feature extraction is still possible, suggesting high robustness of our approach.

In conclusion, Glycan Atlassing provides a comprehensive method for characterizing cellular states by analyzing the nanoscale architecture of the glycocalyx, distinguishing cellular states in different biological systems via the identification of characteristic glycan patterns. The ability to quantify and visualize these glycocalyx signatures at the nanometer scale offers significant potential for advancing our understanding of cellular behavior and developing targeted therapies. Glycan Atlassing is therefore a powerful tool in fundamental glycobiology with the potential to drive innovation in biomedical research and clinical applications.

## Acknowledgements

DMJ, NY, SF, RH, CB, KA, and LM gratefully acknowledge financial support from the Else-Kröner-Fresenius-Stiftung (grant ID 2020_EKEA.91 to LM), the German Research Foundation (DFG, grant ID 529257351 to LM), the Wilhelm-Sander-Stiftung (grant ID 2023.025.1 to LM), and the Max Planck Society. KF and TB gratefully acknowledge financial support from the European Research Council (Synergy Grant 101118729 UNFOLD to KF), the DFG (projects 460333672 CRC1540 EBM and 270949263 GRK2162 to KF) and the Alexander von Humboldt Foundation (Alexander von Humboldt Professorship to KF). We thank the Central Biobank Erlangen [biobank] for excellent support in the conduct of this comprehensive study.

## Methods

### Cell culture

MCF10A/AT cells were cultured in T-25 flasks (Greiner Bio-one, cat# 690175) containing DMEM/F12(Gibco, ref# 21041-025) medium supplemented with 5% horse serum (Gibco, ref# 16050-112), 20 ng/mL epidermal growth factor (EGF) (Gibco, ref# PHG0311), 0.5 μg/mL hydrocortisone (Sigma-Aldrich, ref# H0396), 100 ng/mL cholera toxin (Sigma-Aldrich, cat# C8052-5MG), 10 μg/mL insulin (Sigma-Aldrich, cat# I1882-100MG), and 1% penicillin/streptomycin (Sigma-Aldrich, cat# P0781-100ML). Cells were maintained at 37°C in a humidified atmosphere with 5% CO_2_. Upon reaching 70-80% confluency, cells were washed with 5 ml of DPBS, Ca^2+^/Mg^2+^-free (Gibco, ref# 14190-094) and harvested by enzymatic dissociation by brief incubation with 0.05% trypsin-EDTA solution (Gibco, ref# 2500-054) for 8-10 minutes. Following trypsin neutralization with complete medium, cells were centrifuged at 3000 × rcf for 5 minutes, and the resulting pellet was resuspended in 5 mL of complete growth medium. For experimental assays, cells were seeded into 8-well chamber slides (Ibidi GmbH, ref# 80807-90) at a 1:10 dilution and incubated for 24 hours for prior sample preparation. For the TGFβ treated variants of MCF10A/AT, cells were cultured in media containing 5ng/mL of TGFβ (Bio-Rad, ref# PHP143B) for the following 4 passages. Then the cells were continued in culture using complete media without TGFβ.

A549 cells were cultured in T-25 flasks containing RPMI-1640 medium (Gibco, ref 31870-025) supplemented with 10% fetal bovine serum (FBS) (Gibco, ref# A31605-02), 1% penicillin-streptomycin, and 1% GlutaMAX (Gibco, ref# 35050-38). Cells were maintained at 37°C in a humidified atmosphere with 5% CO2. Upon reaching 70-80% confluency, cells were washed with 5 ml of DPBS, Ca^2+^/Mg^2+^-free, harvested by enzymatic dissociation by brief incubation with 0.05% trypsin-EDTA solution for 8-10 minutes. Following trypsin neutralization with complete medium, cells were centrifuged at 3000 × rcf for 5 minutes, and the resulting pellet was resuspended in 5 mL of complete growth medium. For co-culture experiments, cells were seeded into 8-well chamber slides at a 1:10 dilution and incubated for 24 hours prior to experimental manipulations.

### Isolation and expansion and sample preparation of primary NK cells

NK cells were isolated from peripheral blood using the NK cell isolation kit (Stem Cell Technologies, cat# 19665) following the manufacturer’s instructions, which involves density gradient centrifugation and magnetic bead separation. The isolated NK cells were then expanded using the NK cell expansion kit (Stem Cell Technologies, cat# 100-0711), which includes the base medium (Stem Cell Technologies, cat# 100-0712), supplement (Stem Cell Technologies, cat# 100-0715), and coating material (Stem Cell Technologies, cat# 100-0714). Briefly, a 24-well plate (Stem Cell Technologies, cat# 38044) was coated with the coating material and incubated for 2 hours at room temperature, then rinsed with DPBS, Ca^2+^/Mg^2+^-free. Isolated NK cells were seeded onto the coated plate at a density of 1 x 10^6 cells/mL in ImmunoCult™ NK Cell Expansion Medium and incubated at 37°C and 5% CO2. On Day 3 or 4, an additional expansion medium was added, and the cells were further incubated. On Days 7 and 10/11, the cells were harvested via DPBS, Ca^2+^/Mg^2+^-free, and reseeded onto coated 8 well plates (2 x 10^5 cells/mL) in a fresh expansion medium and incubated for 72 hours to prepare samples for fixation.

### Co-culture of NK cells-A549 cells

A549 cells were seeded in 1:10 concentration onto surface-coated 8-well plates and incubated at 37°C in a humidified atmosphere containing 5% CO_2_ for 24 hours to allow for proper adherence and growth. Separately, previously seeded and expanded NK cells were resuspended for passaging and co-culture experiment, For co-culture, resuspended cells in fresh media subsequently added to the A549 cells in a concentration of 2 x 10^5^ cells/mL. The co-culture was incubated at 37°C in 5% CO_2_ for a duration of 5 minutes, during which NK cells adhered to the A549 cancer cells. Interaction between NK and A549 cells was closely monitored using light microscopy. Once physical interactions between NK and A549 cells were observed, the samples were immediately fixed.

### Preparation of tissue slices

Cryosections were fixed using 4% PFA for 20 minutes and transferred to #1.5 glass slides. Prior to imaging, a chamber (Ibidi sticky slides, ref# 80427) was placed on top of the glass slides and filled with PBS to facilitate staining, imager strand addition and washing steps.

### Metabolic incorporation of Ac4ManNAz and copper-free click chemistry

For metabolic labeling of sialic acids, the MCF10A/T panel was seeded into slides as described above. After 3 hours, the cell culture medium is supplemented with 50 µM Ac_4_ManNAz and incubated for 72 hours. After incubation, cells were washed twice with DPBS, Ca^2+^/Mg^2+^-free and then refreshed with standard growth medium supplemented containing 50 µM DBCO conjugated to docking strand R6 for 2 hours in the incubator to facilitate the click reaction. Then, cells were fixed and stained with lectins.

### Primary neuronal cultures

Hippocampi of E18 Sprague Dawley rats (Transnet YX, SKU# SDEHP) were washed 3x in ice-cold HBSS (ThermoFisher, cat# 14175095) with 1% Pen-Strep (ThermoFisher, cat# 15140122). Tissue was then incubated for 10 min in 0.05% Trypsin EDTA (Gibco, cat# 15400054) at 37°C. Trypsin was removed and tissue washed 10x in pre-warmed HBSS with 1% Pen-Strep. HBSS was replaced by 1 ml of warm Neurobasal Medium (ThermoFisher, cat# 12348017) supplemented with 1% Pen-Strep, 1% GlutaMAX (ThermoFisher, cat# 35050061), and 2% B27 (ThermoFisher, cat# 1750404). Cells were then mechanically dissociated by pipetting up and down about 30 times with a 200 µl pipette. 100,000 – 150,000 cells were seeded on glass-bottomed petri dishes (WPI, cat# FD35-100) coated with 10 µg/ml poly-D-lysine (PDL, Sigma-Aldrich cat# p6407-5mg) and 1 µg/ml laminin (Merck, cat# L2020-1MG) in 2 ml Neurobasal medium supplemented with 1% Pen-Strep, 1% GlutaMAX, and 2% B27. The medium was replaced the next day, and afterwards half of the medium was replaced every 3 days.

### Isolation, activation and preparation of primary human CD4+ T cells

CD4+ T cells were isolated from blood samples of healthy donors. First, peripheral blood mononuclear cells (PBMCs) were collected after Ficoll gradient isolation (Ficoll-Paque® PLUS, VWR). CD4+ T cells were then separated using commercially available CD4+ T cell isolation kit for human samples (Miltenyi), according to manufacturer’s instructions. 0.5·10^6^ CD4+ cells were resuspended in Iscove’s Modified Dulbecco’s Medium (IMDM, Gibco) supplemented with 10% fetal bovine serum (FBS, PanBiotech) and 1% penicillin/streptomycin (P/S, Gibco). To activate the cells, anti-human CD3 (α-CD3, 1 µg/ml, Ultra-LEAF™ Purified anti-human CD3 Antibody, Biolegend) together with anti-human CD28 (α-CD28, 2 µg/ml, Ultra-LEAF™ Purified anti-human CD28 Antibody, Biolegend) and recombinant human IL-2 (20 ng/ml, ImmunoTools) were also provided. The cells were cultured for three days at 37°C and 5% CO_2_. For control, non-stimulated and freshly isolated CD4+ T cells from same donor were used. Samples were then fixed using 4% paraformaldehyde (PFA) for 10 minutes at room temperature and washed three times with PBS before lectin labelling.

### Isolation, stimulation and preparation of primary human neutrophils

Human neutrophils were isolated using a commercially available kit (MACSxpress® Whole Blood Neutrophil Isolation Kit, Miltenyi), according to the manufacturer’s instructions. 10^6^ neutrophils were then prepared in RPMI1640 medium (Gibco) supplemented with 10% fetal bovine serum (FBS, PanBiotech) and 1% penicillin/streptomycin (P/S, Gibco). For stimulation conditions, recombinant human tumor necrosis factor-alpha (rh TNF-α, 100 ng/ml, Immunotools) was also added and the cells were cultured for two hours at 37°C and 5% CO2. Non-stimulated neutrophils were kept in the same conditions for culture, but without TNF-α. Samples were fixed using 4% PFA for 10 minutes at room temperature and washed three times with PBS before lectin labelling.

### Sample fixation

Samples were washed with DPBS, Ca^2+^/Mg^2+^ (Gibco, ref# 14040-091) 3 times. Then 4% paraformaldehyde, diluted from a 16% stock solution (Thermo Scientific, ref.no: 28908) to 4% working solution via DPBS, Ca^2+^/Mg^2+^-free was added to wells and cells were incubated at room temperature for 15 minutes. Then, cells were washed three times with DPBS, Ca^2+^/Mg^2+^-free. After fixation, cells were permeabilized with 0.1% Triton-X (Alfa Aesar, cat# A16046) for 10 minutes at room temperature, followed by four DPBS, Ca^2+^/Mg^2+^-free washing steps.

### Lectin labeling

A lectin cocktail (2.5μg/mL of each lectin) prepared in 1X tris buffer (Fisher Bioreagents, ref# M-15836) was applied to the cells at room temperature for 30 minutes, allowing specific binding to glycan targets on the cellular surface. Following incubation, cells were washed three times with DPBS, Ca^2+^/Mg^2+^-free. For MCF10A panel cells, a pre-permeabilization is performed using 0.1% Triton-X 100 for 10 minutes at room temperature, followed by four washing steps with DPBS, Ca^2+^/Mg^2+^-free and incubation with lectins as above. Prior to imaging, another permeabilization step using 0.1% Triton-X 100 was performed for 10 minutes at room temperature, followed by four washing steps with DPBS, Ca^2+^/Mg^2+^-free. For primary neurons, a permeabilization step using 0.1% Triton-X 100 was performed for 10 minutes at room temperature, followed by four washing steps with DPBS, Ca^2+^/Mg^2+^-free prior to imaging. For primary immune cells and tissue sections, samples were permeabilized with 0.2% Triton-X 100 for 10 minutes at room temperature, followed by four washing steps with DPBS, Ca^2+^/Mg^2+^.

### Optical setup

DNA-PAINT imaging was carried out on an inverted microscope (Nikon Instruments, Eclipse Ti2) with the Perfect Focus System. Objective-based Total Internal Reflection Fluorescence (TIRF) mode was used, employing a high NA objective (Nikon Instruments, Apo SR TIRF×100, NA 1.49, Oil) and the Nikon TIRF module. A 560-nm laser (MPB Communications, 1 W) was used for excitation and coupled into the microscope via the TIRF module. The power of the laser beam was controlled in free space using a filter wheel (Thorlabs, FW212CNEB). The laser beam was passed through a cleanup filter (Chroma Technology, ZET561/10) and coupled into the microscope objective using a beam splitter (Chroma Technology, ZT561RDC). Fluorescence was spectrally filtered with an emission filter (Chroma Technology, ET600/50m, and ET575LP) and imaged on an sCMOS camera (Hamamatsu Orca Fusion) without further magnification, resulting in an effective pixel size of 130 nm after 2×2 binning. TIRF illumination was used for all measurements with a laser power of approx. 33 mW above the objectiuve. The central 1152×1152 pixels (576×576 after binning) of the camera were used as the region of interest. Raw microscopy data was acquired using μManager (Version 2.0.3).

### DNA sequences

Docking and imager strand sequences are previously optimized for orthogonal binding specificity. The sequences used for DNA-PAINT were as follows:

R1 (WGA) with 5×R1 sequence TCCTCCTCCTCCTCCTCCT

R2 (SNA) with 5×R2 sequence ACCACCACCACCACCA

R3 (PHA-L) with 7×R3 sequence CTCTCTCTCTCTCTCTCTC

R4 (AAL) with 7×R4 sequence ACACACACACACACA

R5 (PSA) with 5×R5 sequence CTTCTTCTTCTTCTT

R6 (DBCO) with 5×R6 sequence AACAACAACAACAACAA.

All DNA oligonucleotides were HPLC-purified and obtained from Metabion.

### Imager Strand Preparation for DNA-PAINT Imaging

Imaging buffer was prepared by combining 50 ml of DPBS, Ca^2+^/Mg^2+^-free with 0.0146 g ethylenediaminetetraacetic acid (EDTA) (PanReac, cat# 60-00-4), 1.461 g sodium chloride (NaCl) (Alfa Aesar, cat# A12313), and 10 µl TWEEN-20 (MP Biomedicals, cat# 103368) in a 50-ml falcon tube. All components were thoroughly mixed until completely dissolved. The prepared buffer was stored at 4°C until use in subsequent imager strand preparation. Imaging strands for DNA-PAINT were prepared by diluting 1 µM stock solutions in the buffer, the optimal imager concentration to achieve sparse blinking ranged from 0.075 - 0.5 nM, adjusted for sample types and targets, focusing on sparse single-molecule signals. Samples were incubated with a 1:3 dilution of 90 nm gold nanoparticles (Absource, cat# CG-90-20), which were used as fiducial markers for drift correction and channel alignment.

### Multiplexed DNA-PAINT

DNA-PAINT imaging was conducted via six subsequent imaging rounds with only one imager type in each round. For each target, 20,000 frames of single molecule blinks are captured with a frame time of 100 milliseconds. A laser power of approx. 33mW at the objective was used. Between imaging rounds, thorough washing of at least four times with DPBS, Ca^2+^/Mg^2+^-free was performed, ensuring no residual signal from the previous imager solution remained before introducing the next imager solution.

### Postprocessing

Raw fluorescence data were reconstructed using the Picasso software package,^27^ version 0.7.4. For identifying distinct blinks in each frame, an intensity threshold of 5000 was used. Drift correction was performed using redundant cross correlation, followed by precise fiducial-based drift correction with gold nanoparticles. The six channels were aligned using gold nanoparticles as fiducial markers. Region of interest is segmented using the polygon pick function from Picasso. All downstream analysis after segmentation until calculation of cluster centers was performed using a custom version of Picasso in python. An estimate of experimental localization precision is calculated for each individual channel of images using nearest neighbor-based analysis (NeNA).^51^ For clustering, a radius of 2 times the experimental NeNA precision was used. The minimum number of localizations inside a cluster was two. Time signatures of blinking events inside each cluster were analyzed to account for unspecific sticking events. If for a single cluster a time bin of 200 frames (1% of the total length of the stack) contains more than 90% of all the events in that cluster, the respective cluster is rejected. Thus, single and/or atypically extended events are rejected as false localization. Cluster centers are calculated and are used as the location of glycan targets.

For further analysis, two approaches were used. First, we calculated the NN distances across all glycan localizations, both within channels and between different channels. Histograms of the distances to the first NN were plotted and the peak of the distributions were collected in a 5×5 matrix (6×6 for datasets with DBCO). Second, we performed a classification of lectin binding sites that are frequently occurring in close spatial proximity using a cutoff radius of 5 nm. Each observed combination of two or more lectin binding sites was assigned a different class. The distribution of classes per square microns was plotted and the cellular location of these classes are mapped. To handle this multidimensional data, we performed standard linear dimensionality reduction using PCA in python.

