## Supplemental Information for "Glycan Atlassing: Nanoscale analysis of glycocalyx architecture enables functional tracing of cell state"

Electronic Supplementary Information

**Table of contents**

Figure S1: Additional reconstructions across sample types p. 2
Figure S2. Quality metrics p. 3
Figure S3: Further data on MCF10A panel p. 6
Figure S4: Further data on primary neurons p. 10
Figure S5: Further data NK cells p. 11
Figure S6: Further data on tissue p. 12

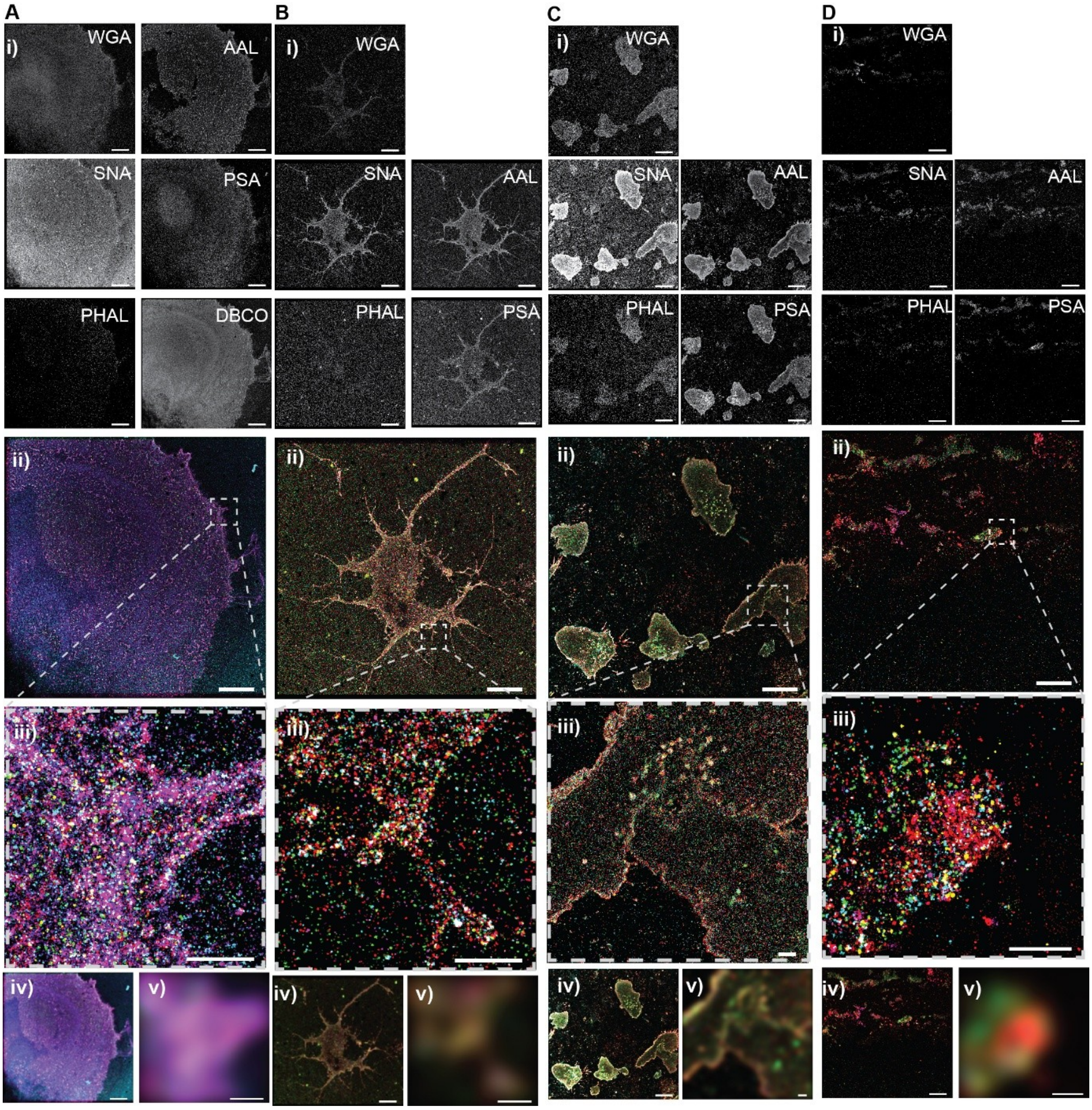


**Figure S1: Additional reconstructions across sample types.** A-D) Individual channels depicted in grayscale (i), merged channels (ii) using the following color code: WGA – magenta, SNA – cyan, PHA-L – yellow, AAL – red, PSA – green, DBCO – purple. iii) Zoom-in showing intricate details resolved. iv) Diffraction-limited representation of the whole field of view. v) Diffraction-limited zoom-in corresponding to (iii). A) MCF10AT, B) Primary neuron, C) Immune cells, D) Tissue. Scale bars: 10 µm for full field of views and 1 µm for zoom-ins.

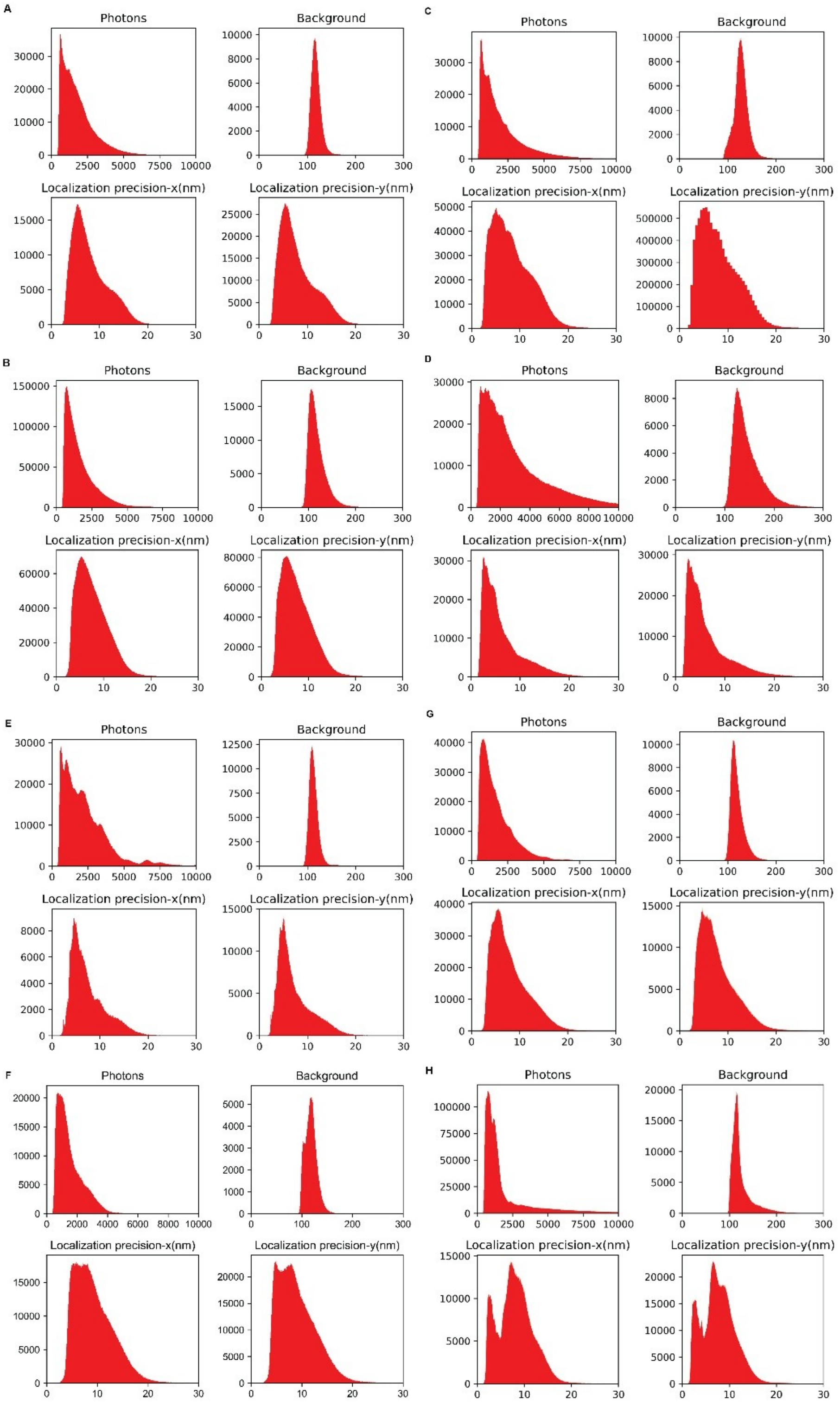


**Figure S2: Quality metrics.** Signal photon count, background, localization precision in x and y of a representative channel (AAL) for **A)** MCF10A **B)** MCF10A+TGFβ **C)** MCF10AT **D)** MCF10AT+TGFβ **E)** Neurons **F)** Tissue sections **G)** Stimulated NK cells **H)** Non stimulated NK cells **I)** Non stimulated CD4+ cells **J)** Stimulated neutrophils **K)** Non stimulated neutrophils. **L)** Localization precision across channels and across the whole dataset used in the study (see plot titles). Localization precision are given as NeNA precisions.

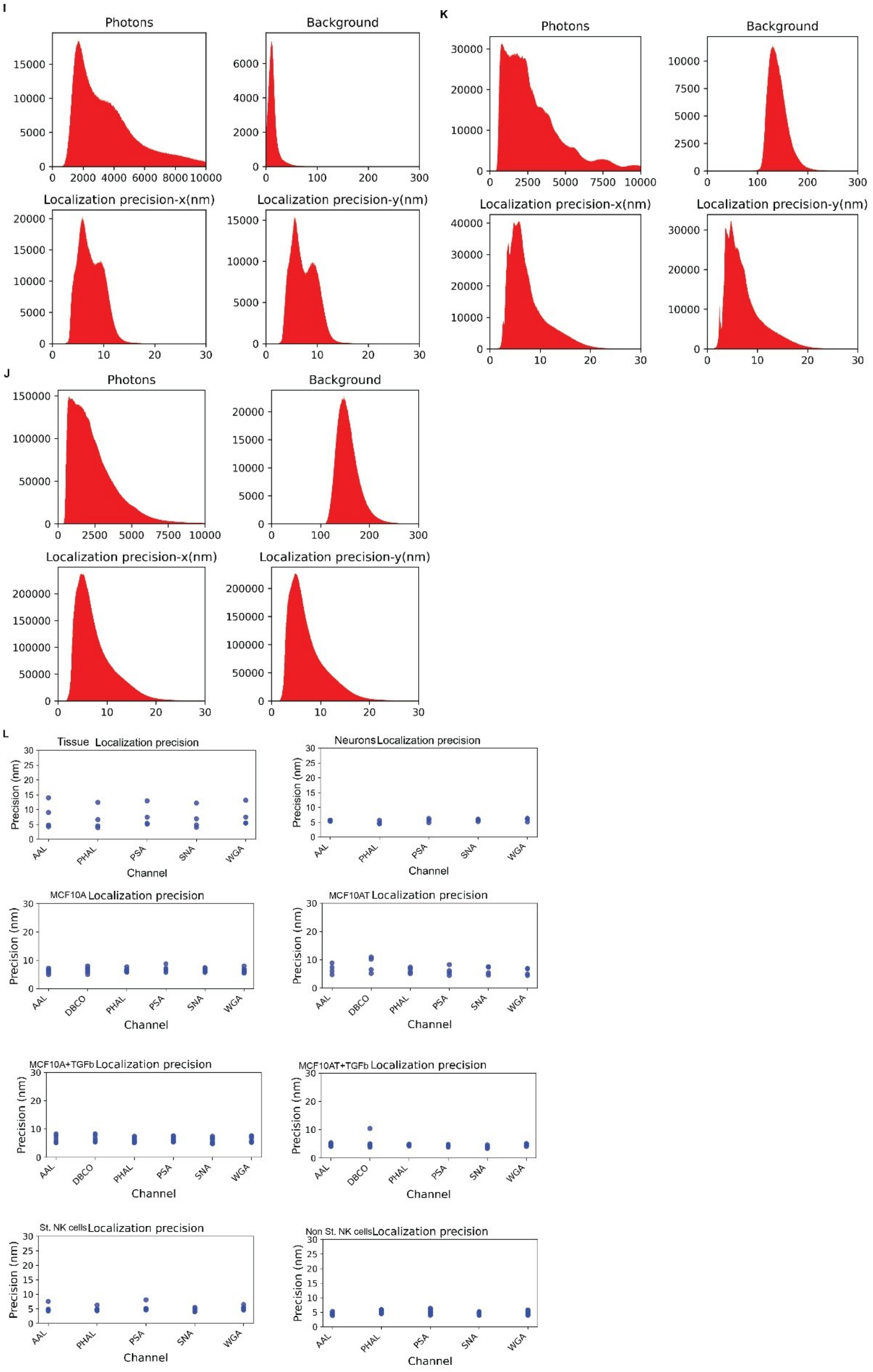


**Figure S2: Quality metrics continued.**

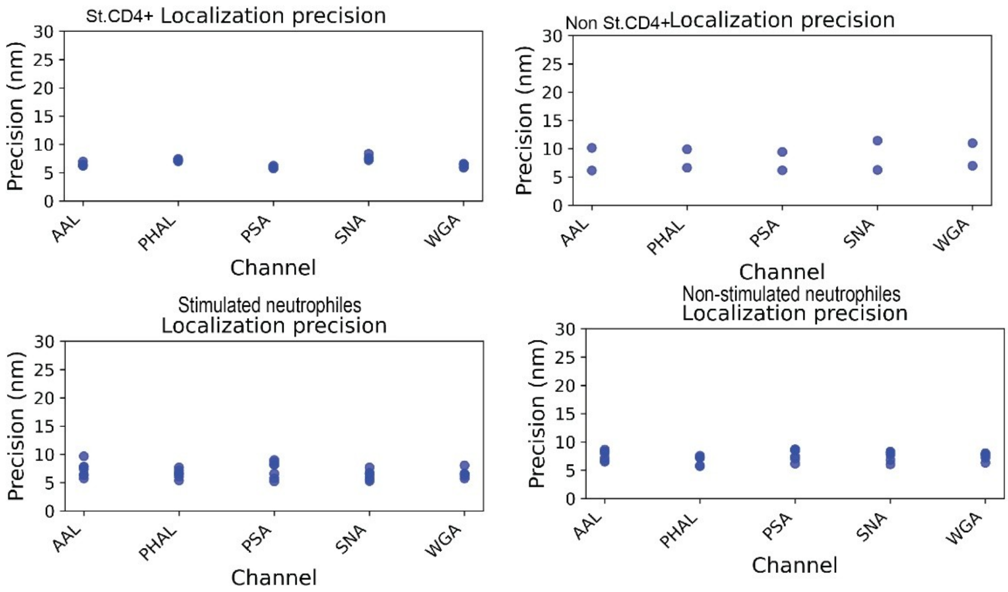


**Figure S2: Quality metrics continued.**

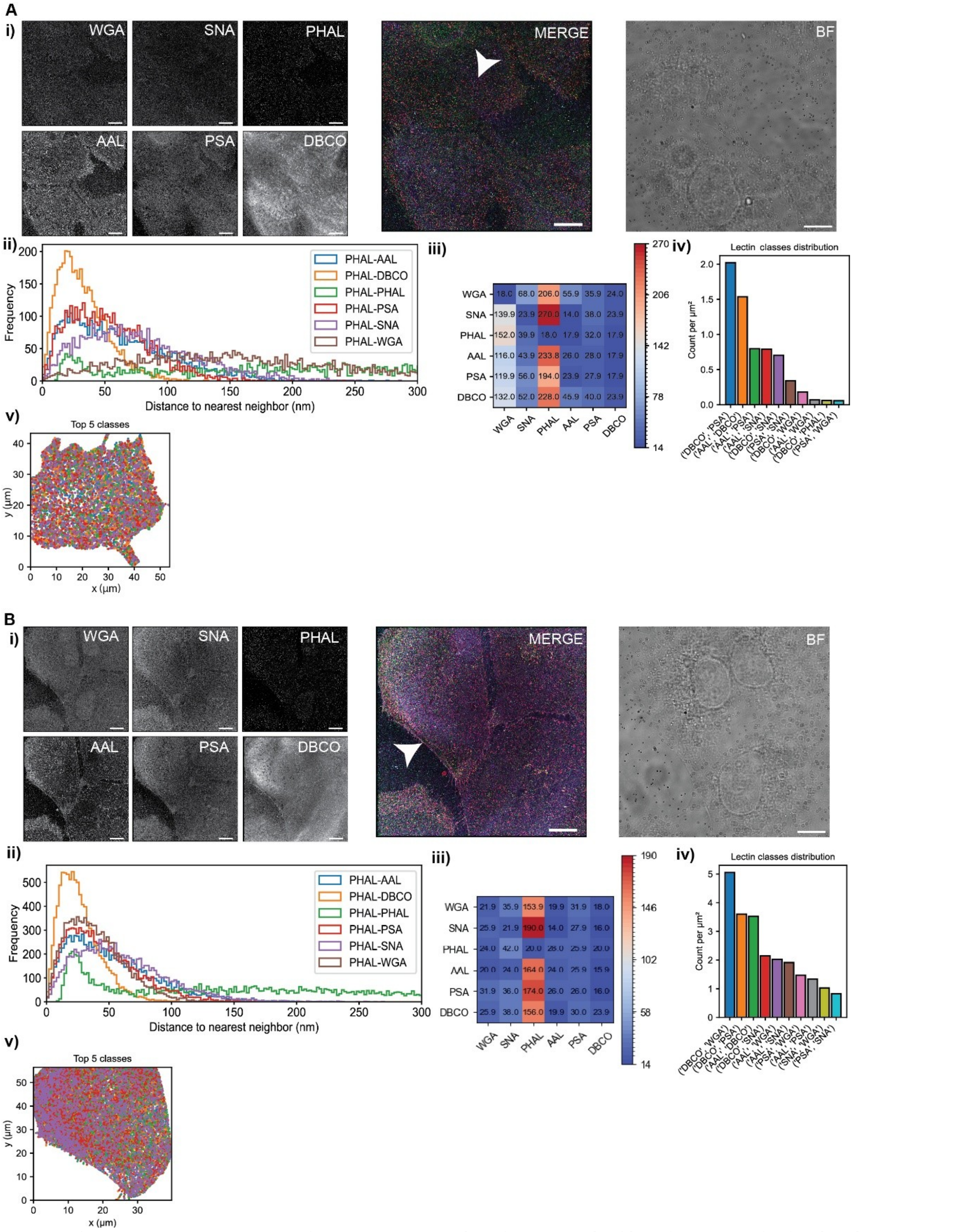


**Figure S3: Further data on MCF10A panel. A)** MCF10A; **B), C)** MCF10A+TGFβ **D), E)** MCF10AT **F), G)** MCF10AT+TGFβ.

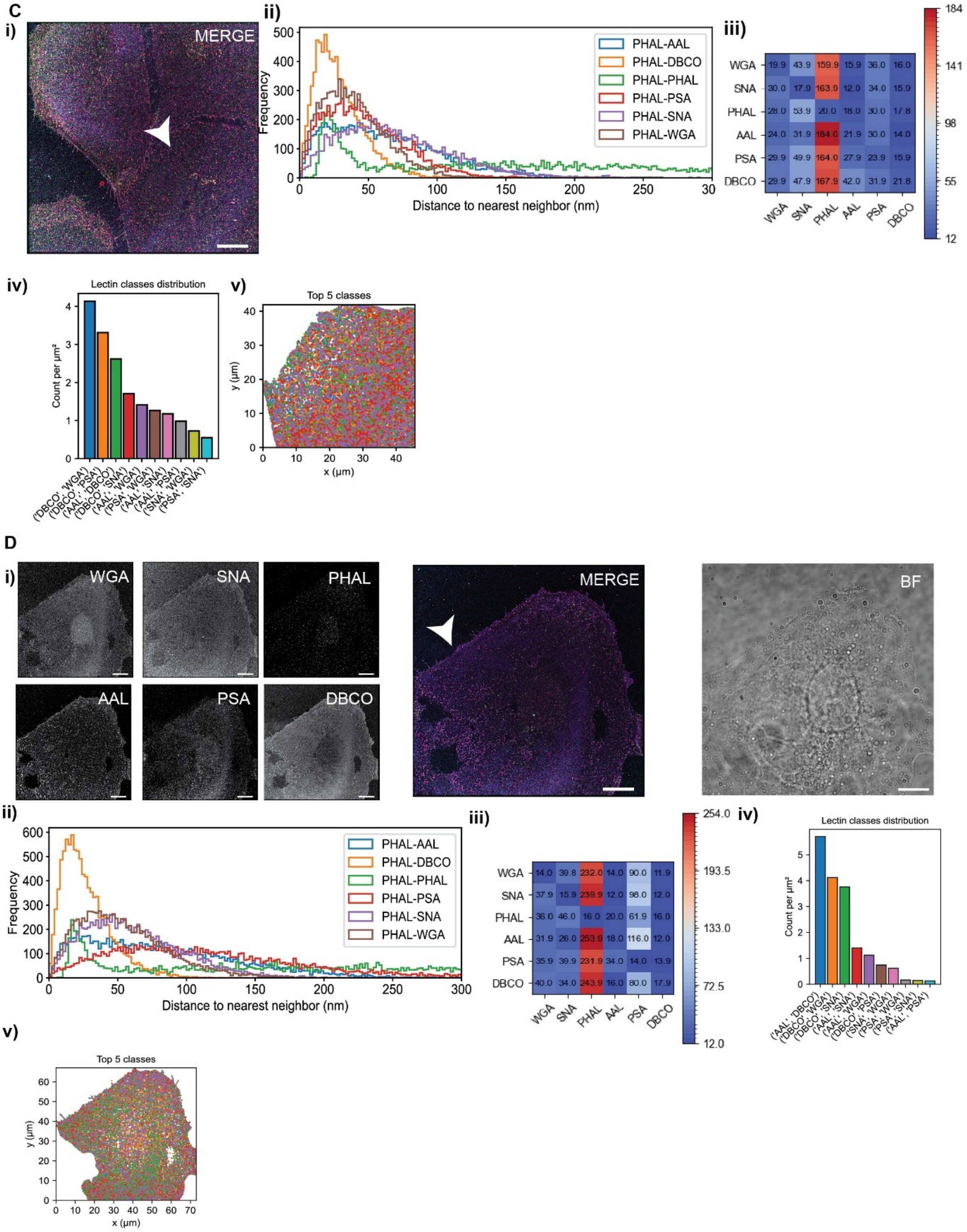


**Figure S3: Further data on MCF10A panel continued.**

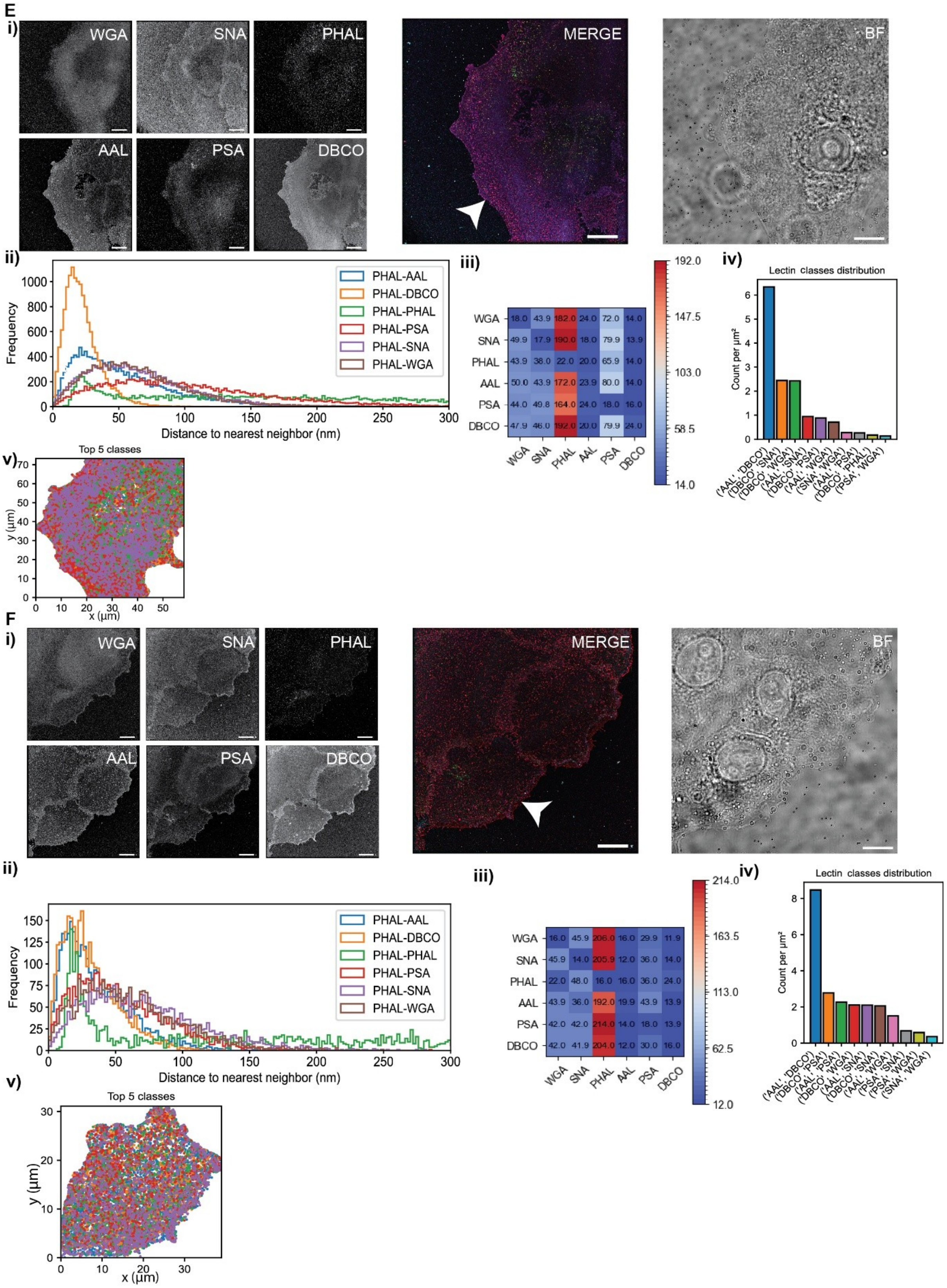


**Figure S3: Further data on MCF10A panel continued.**

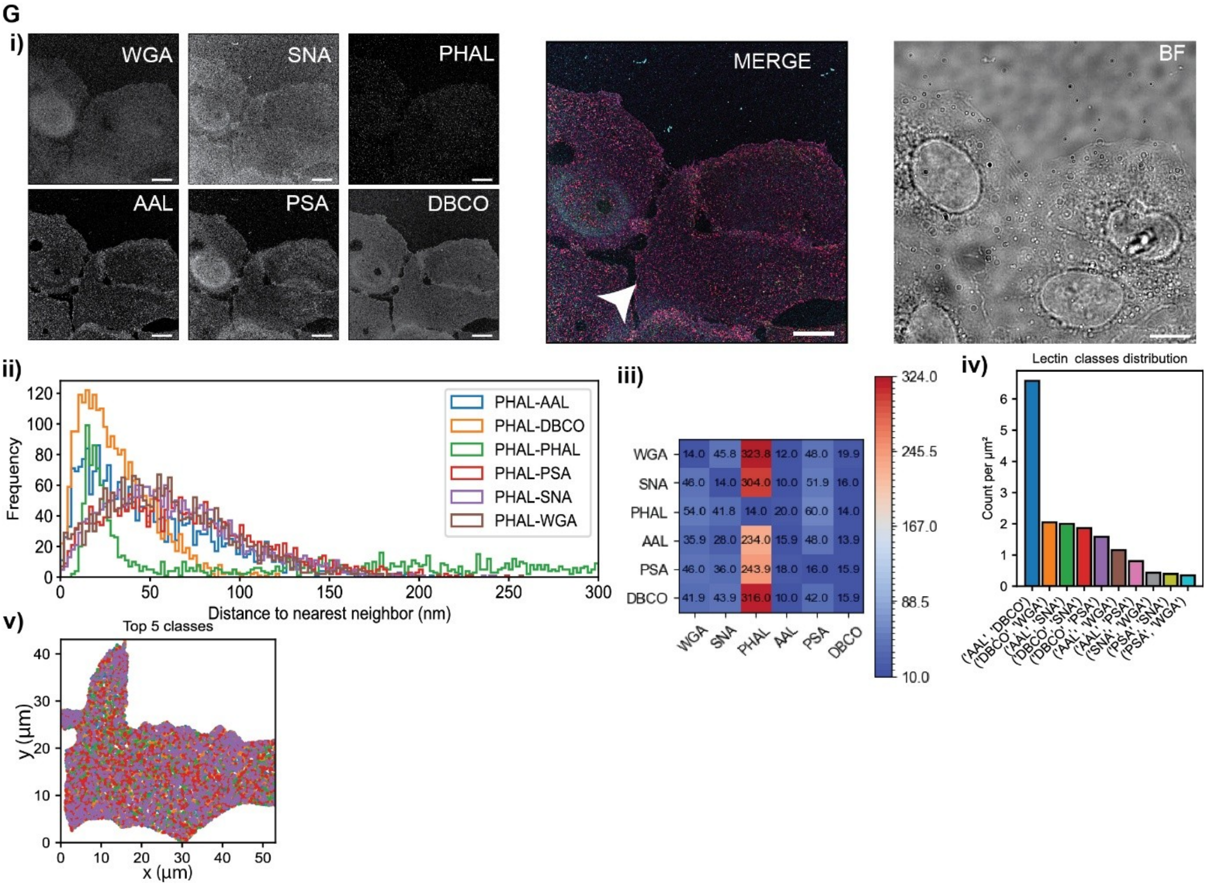


**Figure S3: Further data on MCF10A panel continued.**

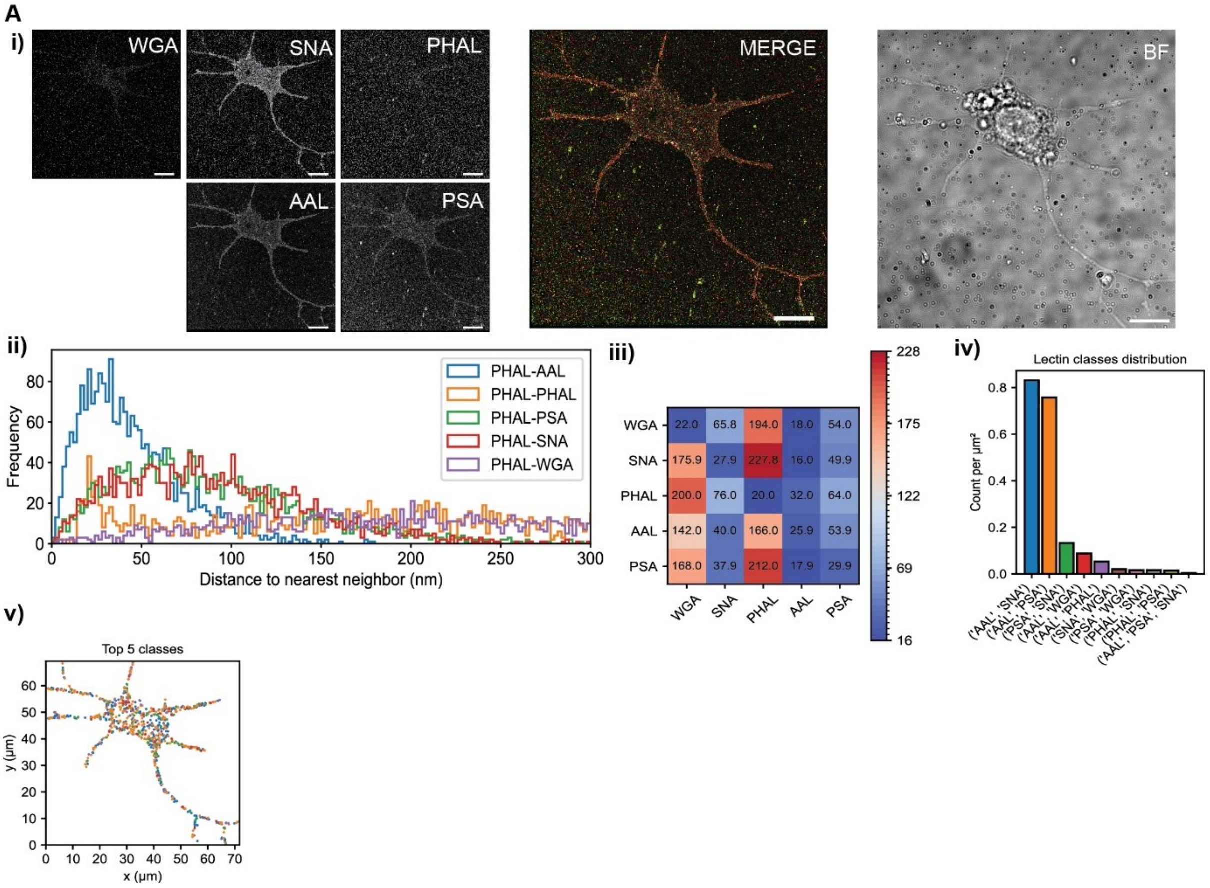


**Figure S4: Further data on primary neurons.**

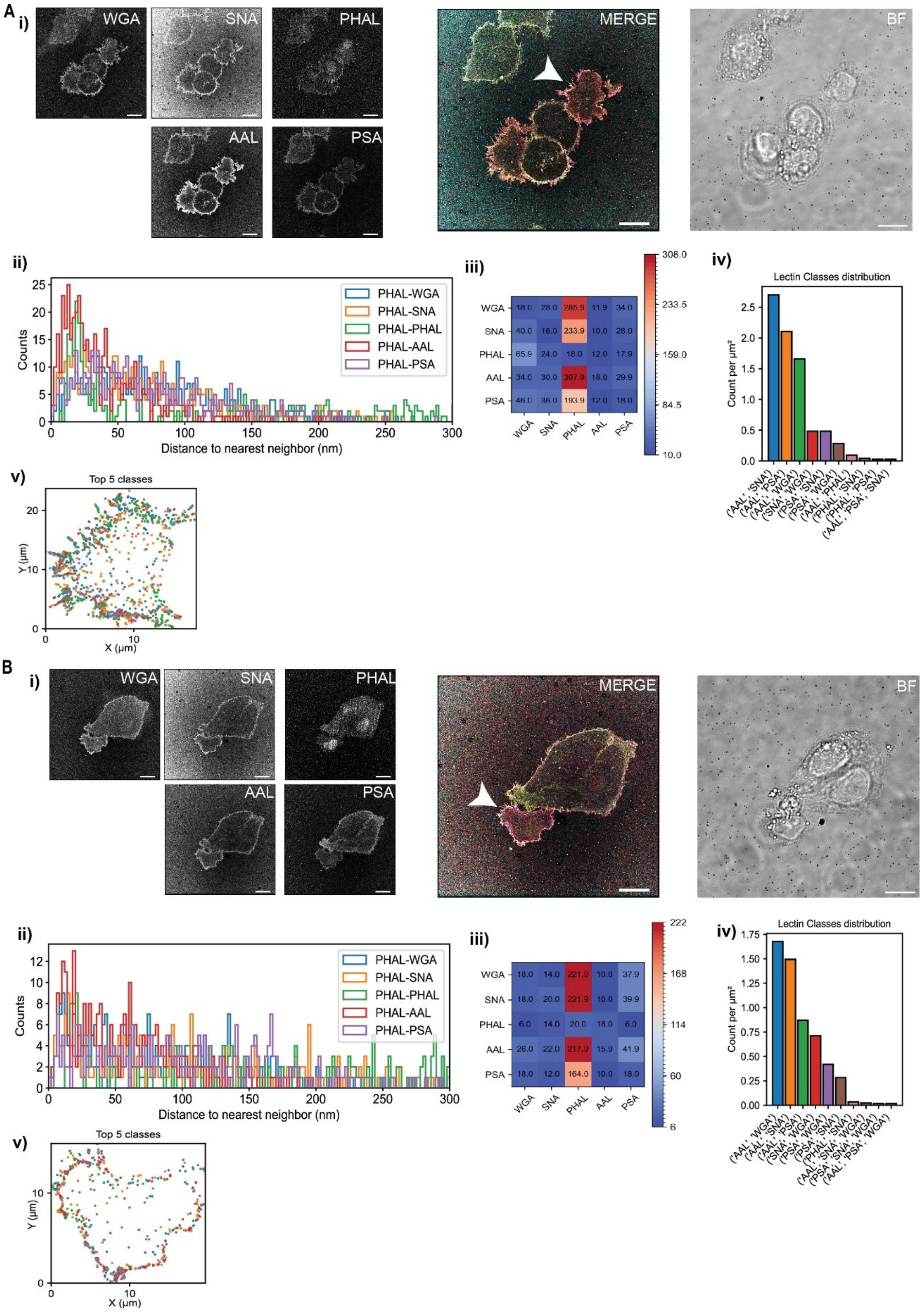


**Figure S5: Further data on NK cells. A), B)** Two representative fields of view, showing NK cells in co-culture with A549 cells.

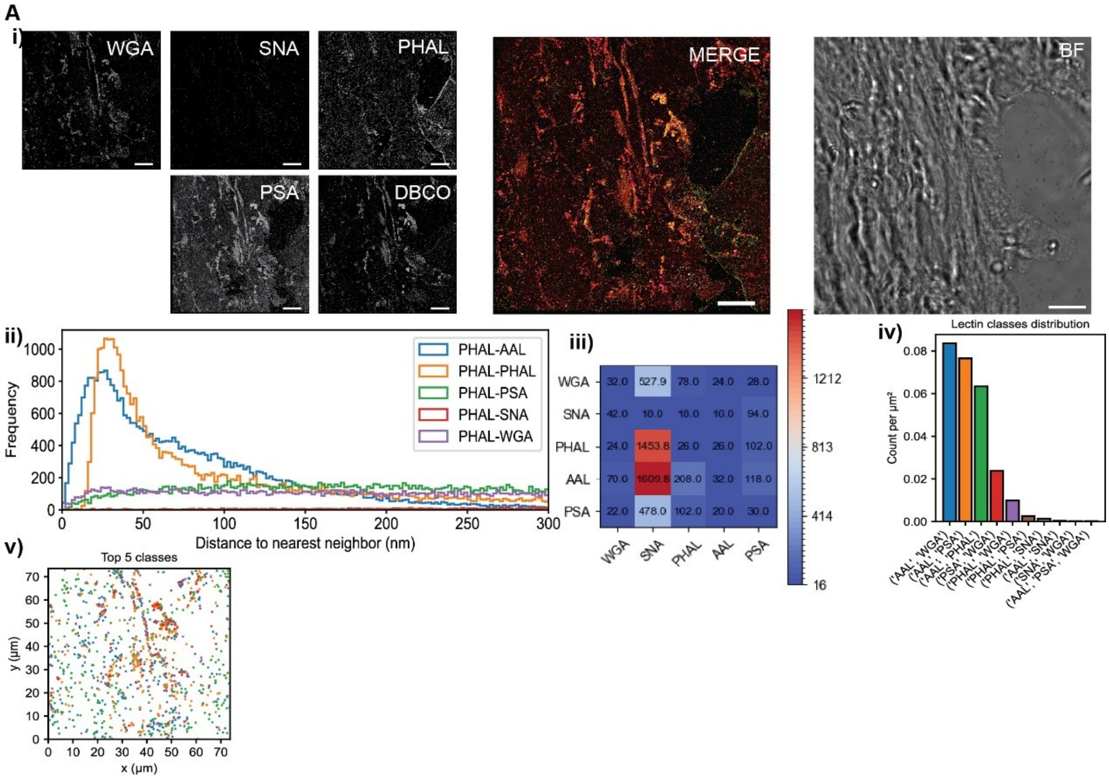


**Figure S6: Further data on tissue.**
